# Predicting Enzyme pH Optima from Structure Using Equivariant Graph Neural Networks

**DOI:** 10.64898/2026.01.18.700076

**Authors:** Rajarshi SinhaRoy, Christian Clauß, Ivan Ivanikov, Georg Künze

## Abstract

Enzyme activity and stability are strongly modulated by pH, making the catalytic pH optimum (pH _opt_) a key parameter in enzyme development and biotechnological applications. Experimental determination of pH _opt_ is, however, labor-intensive and time-consuming, motivating the development of accurate computational prediction methods.

Here, we introduce pHoptNN, an *E*(*n*)-equivariant graph neural network designed to predict enzyme pH _opt_ directly from three-dimensional protein structures. pHoptNN was trained on a curated dataset comprising nearly 12,000 enzymes with experimentally determined pH _opt_ values and high-confidence structural models obtained from the Protein Data Bank and AlphaFold3. The model represents enzymes as atomic-level molecular graphs, integrating structural, chemical, and electrostatic features. Model development was guided by extensive hyperparameter optimization using genetic and Bayesian search strategies.

On a held-out test set, pHoptNN achieved a root-mean-square error (RMSE) of 0.588 pH units, substantially outperforming the sequence-based method EpHod (RMSE = 0.879). Moreover, pHoptNN maintains robust predictive performance across different enzyme classes and pH ranges. These results demonstrate the utility of structure-based equivariant deep learning for enzyme pH _opt_ prediction and highlight the potential of pHoptNN to accelerate enzyme discovery and engineering workflows.

## Introduction

Enzymes are highly efficient biological catalysts that accelerate the attainment of chemical equilibrium without altering its position, thereby enabling biochemical reactions to proceed with exceptional specificity. According to their reaction chemistry and substrate type, enzymes are categorized into seven major enzyme commission (EC) classes. ^1^ Their catalytic activity is governed by the precise spatial arrangement of residues in the active site and is modulated by intrinsic factors such as amino acid composition and post-translational modifications, as well as by extrinsic physicochemical conditions including substrate concentration, temperature, ionic strength, and pH. Among these parameters, pH plays a particularly critical role. It influences the protonation states of catalytic and regulatory residues, thereby affecting substrate binding, transition-state stabilization, and overall catalytic efficiency. In addition, pH can alter tertiary and quaternary structural stability by perturbing hydrogen-bond networks and hydrophobic packing. Deviations from the optimal pH (pH _opt_), i.e., the pH at which an enzyme exhibits maximal catalytic activity, can lead to reversible or even irreversible denaturation.^2^ As a result, pH dependence is a key determinant of enzymatic performance in natural environments and an essential consideration in enzyme engineering for biotechnology, pharmaceutical synthesis, and diagnostic applications.

Despite its importance, experimentally determined pH _opt_ values remain unavailable for the majority of known enzymes. Major biological repositories such as UniProt^3^ and BRENDA^4^ contain extensive annotations on protein sequence and function, yet pH-related information is sparse and heterogeneous, frequently derived from diverse literature sources with variable assay conditions. Existing measurements are heavily biased toward well-studied model enzymes, leaving much of the protein sequence and structure landscape uncharacterized with respect to catalytic pH preference. This information gap persists despite rapid growth in protein sequence and structural data. As of August 2025, UniProt ^3^ contains more than 573,000 reviewed protein sequences, including over 285,000 enzymes, while the Protein Data Bank (PDB)^5^ hosts over 230,000 experimentally resolved structures. These expansions have been driven by advances in high-throughput sequencing technologies^6,7^ and modern structural biology techniques such as cryo-electron microscopy, NMR spectroscopy, and X-ray crystallography. However, the accumulation of experimentally determined biochemical property annotations, including pH _opt_, has not kept pace with this growth.

Computational approaches provide a promising avenue to bridge this annotation gap by predicting biochemical properties from existing sequence and structural data at scale. Several sequence-based methods for catalytic pH prediction have been introduced. For example, EpHod,^8^ integrates embeddings from large-scale protein language models (ESM-1v)^9^ into a residual regression architecture trained on nearly two million sequences with environmental pH annotations, and subsequently fine-tuned on experimentally validated pH _opt_ values. Other sequence-based models, such as that of Yan et al.^10^ employ physicochemical descriptors and amino acid composition to capture predictive trends. More recent sequence-based methods have further improved performance by combining protein language model embeddings with deep learning and retrieval-based strategies. Seq2pHopt^11^ extends the Seq2Topt^11^ framework to enzyme pHopt prediction, showing that sequence-only deep learning models can achieve strong predictive performance on enzyme property prediction tasks. CatOpt^12^ extends the concept by integrating ESM-2 embeddings with multi-scale convolutional neural networks, multi-head self-attention, and residual dense blocks, which improved both prediction accuracy and interpretability through residue-level attention analysis. Retrieval-enhanced approaches have also emerged. VENUS-DREAM ^13^ formulates pHopt prediction as a few-shot meta-learning task using ESM-2 embeddings, *k*-nearest-neighbor retrieval, and Reptile-based adaptation. Moreover, rather than analyzing enzyme sequences in isolation, OpHReda^14^ employs a retrieval-based approach that jointly models the target sequence alongside embeddings of similar known enzymes, thereby achieving significantly higher predictive accuracy. While these methods demonstrate the utility of sequence features, they inherently lack explicit three-dimensional structural information. Because protonation equilibria, electrostatic interactions, active-site geometry, and residue packing critically shape an enzyme’s pH dependence, models that do not represent spatial context and atomic interactions cannot fully account for the mechanistic determinants of pH _opt_.

Structure-based approaches have shown considerable success in related prediction tasks. For example, ThermoMPNN, ^15^ which integrates three-dimensional structural information into model architectures, accurately predicts thermodynamic stability changes (ΔΔ*G*) of protein mutations. These results underscore the potential of structure-aware models to capture the physicochemical principles underlying complex biochemical properties. Graph neural networks (GNNs) are particularly well-suited for protein modeling because proteins can be naturally represented as graphs, where atoms constitute nodes and covalent or spatial interactions form edges. GNNs preserve both connectivity and geometric information and have been widely applied across computational biology, including protein–protein interaction prediction,^16,17^ drug-target affinity modeling,^18–20^ and molecular property estimation.^21,22^ In protein–protein interaction modeling, geometric equivariance has become an increasingly important inductive bias for capturing the spatial nature of molecular systems. SE(3)-equivariant models such as EquiDock demonstrate how rigid protein docking can be performed by directly learning equivariant transformations between unbound protein structures, avoiding exhaustive conformational search while preserving rotational and translational symmetries.^23^ Beyond docking, recent protein–protein interaction site (PPIS) predictors, including MEG-PPIS and EquiPPIS, apply *E*(3)-equivariant message passing over protein structural graphs to more accurately model residue-level interactions, leading to improved performance in interaction site identification.^24,25^

Related ideas have also been influential in drug–target affinity prediction, where graph-based representations have largely replaced sequence-only approaches. GraphDTA is a representative example that models drug molecules as graphs and uses GNNs to predict binding affinities, achieving strong performance across multiple benchmark datasets.^26^ More broadly, equivariant learning has played a central role in molecular property prediction and quantum chemistry. SchNet introduced continuous-filter convolutions on atomic graphs to learn quantum interactions in a data-driven manner,^27^ while PaiNN further incorporated rotational equivariance through vector-valued message passing, enabling accurate prediction of both scalar and tensorial properties.^28^ *E*(3)-equivariant models such as NequIP have since pushed performance further by explicitly preserving physical symmetries when learning interatomic potentials, underscoring the importance of equivariance for data efficiency and physical consistency in molecular modeling.^29^

Equivariant GNNs, like the *E*(*n*)-equivariant graph neural network (*E*(*n*)-EGNN)^30^ used here, extend these capabilities by incorporating rotational and translational symmetry, ensuring that predictions remain independent of molecular orientation. These architectures have demonstrated state-of-the-art performance on datasets like QM9,^31^ which include molecules with highly complex geometric and energetic properties.

Here, we introduce a structure-based framework for predicting catalytic pH _opt_ values that combines atomistic protein representations with an equivariant message-passing architecture. Our approach advances beyond previous work in three major ways. First, proteins are represented as atom-level graphs featuring hybrid edge descriptors that integrate covalent bonding, ring membership, and distance-based interactions to retain both chemical integrity and spatial neighborhood information. Second, we employ an *E*(*n*)-equivariant GNN that jointly updates node features and atomic coordinates, enabling the model to learn structural patterns from both local chemical environments and global protein geometry. This architecture can optionally incorporate attention mechanisms to highlight structural interactions most relevant for prediction, thereby aiding interpretability. Third, we address the pro-nounced imbalance in the pH _opt_ distribution by applying label distribution smoothing and inverse-frequency reweighting, which improve model robustness in underrepresented acidic and alkaline regions.

To enable large-scale training of structure-based models, we assembled EnzyBase12k, a comprehensive dataset comprising nearly 12,000 enzymes with experimentally determined pH _opt_ values and high-quality three-dimensional structures sourced from the Protein Data Bank^5^ and AlphaFold. ^32,33^ By integrating curated functional annotations with reliable structural models, EnzyBase12k provides broad coverage across enzyme classes and pH ranges and serves as a robust foundation for training structure-based predictive models.

Building on this dataset, we developed a structure-based prediction framework that integrates atomistic graph representations with an *E*(*n*)-equivariant message-passing architecture (Figure 1). The workflow comprises four major components: (i) preprocessing of protein structures into detailed atom-level graphs; (ii) equivariant message passing using EGNN layers that jointly update node embeddings and coordinates; (iii) training with hyperparameter-optimized, imbalance-aware loss functions to address the skewed pH _opt_ distribution; and (iv) inference using a graph-level decoder that produces pH _opt_ predictions along with attention-based relevance scores for interpretability. The resulting model, pHoptNN, achieves a test-set RMSE of 0.588, substantially outperforming the sequence-based benchmark EpHod ^8^ (RMSE 0.879), and exhibits strong generalization across enzyme classes and pH regimes. Beyond predictive accuracy, the model identifies structural features associated with pH _opt_ variation, including catalytically relevant residues and solvent-exposed regions involved in electrostatic stabilization, thereby offering mechanistic insight into the molecular basis of enzymatic pH adaptation.

**Figure 1:**
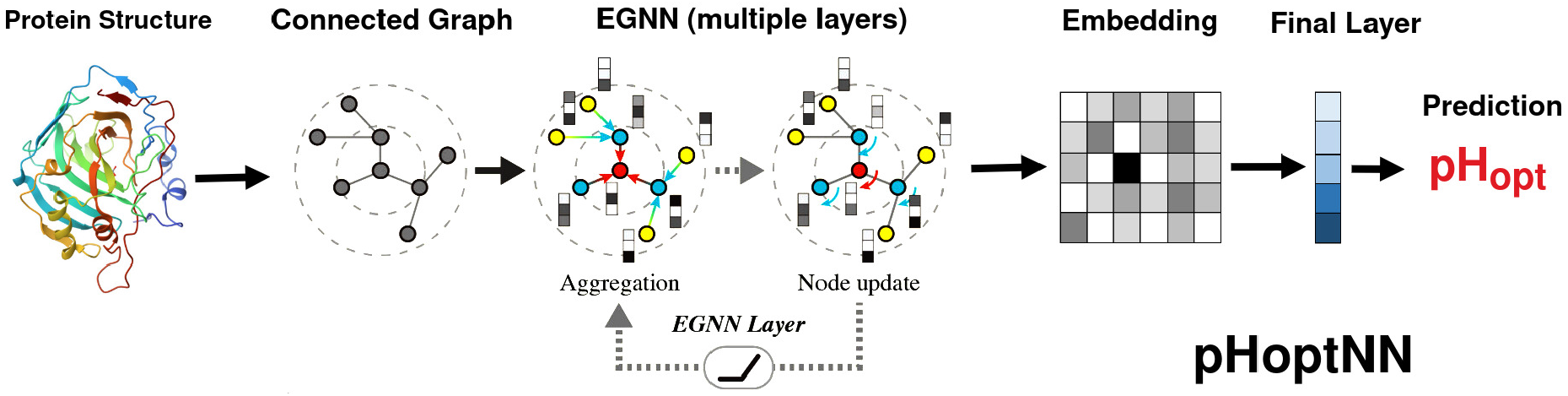
Schematic overview of the pHoptNN workflow for enzyme pH _opt_ prediction. Three-dimensional protein structures are represented as connected atomic graphs. Node and edge features are iteratively updated through stacked *E*(*n*)-equivariant graph neural network (EGNN) layers, enabling the model to capture spatial, structural, and physicochemical relationships. The final layer maps the learned graph embeddings to a predicted catalytic pH optimum.

## Methods

### Dataset Curation and Preprocessing

The construction of the EnzyBase12k dataset followed a multi-stage curation workflow as shown in Figure 2. From 96,955 BRENDA^4^ records accessible in release 2023.1 (version 1.1.0), we first extracted 11,001 enzymes for which both a catalytic pH optimum and a corresponding UniProt^3^ identifier were available. Entries associated with deprecated or inaccessible UniProt identifiers were removed, and duplicate mappings arising from multi-functional enzymes were consolidated, yielding 9,766 unique enzyme–pH pairs. For enzymes with multiple reported pH optima, we retained the mean value when the reported range was ≤ 1 pH unit; otherwise, the record was excluded. This filtering step resulted in 9,537 high-confidence entries derived from BRENDA.

**Figure 2:**
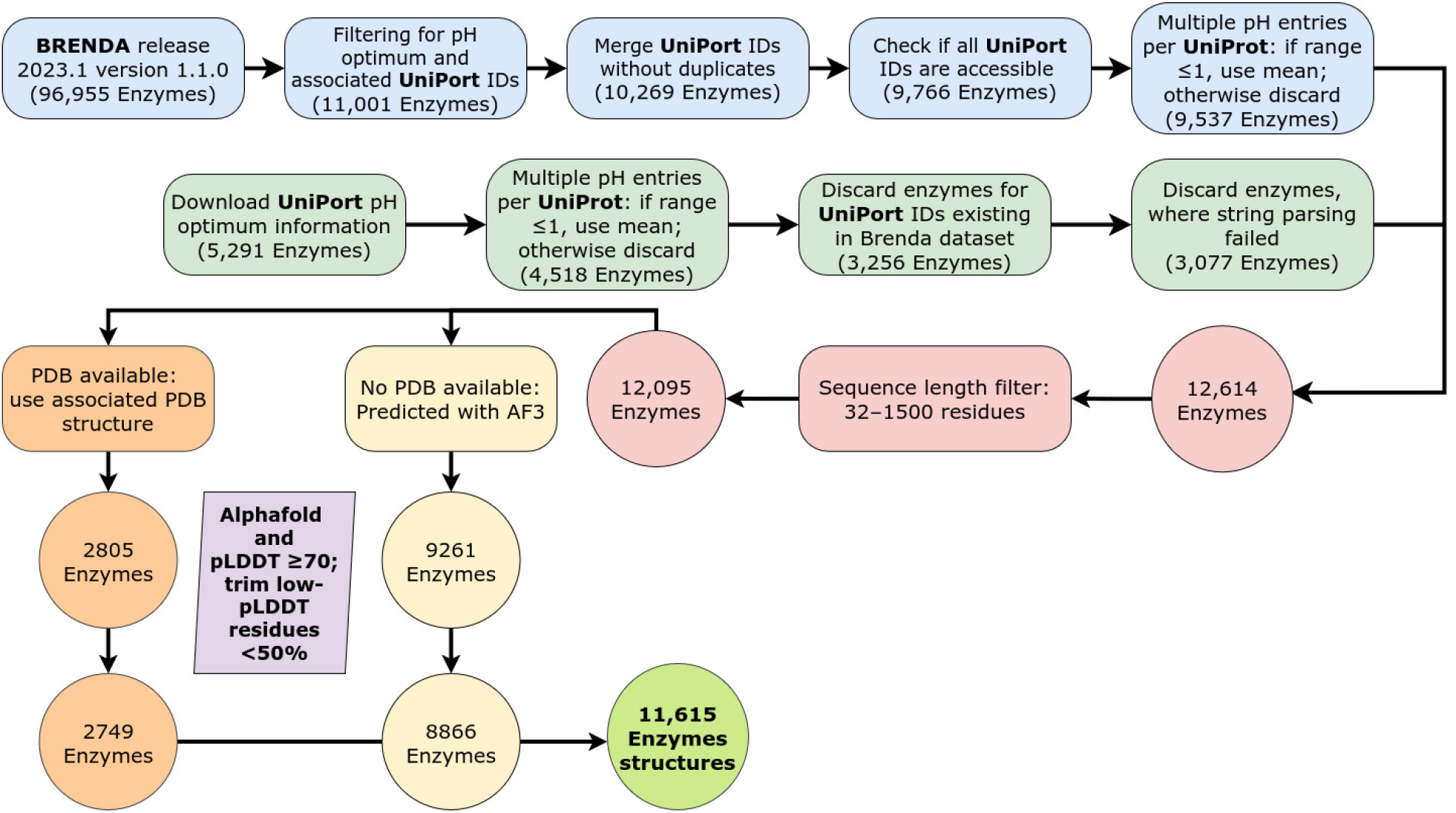
Curation workflow of the EnzyBase12k dataset. (See also Supplementary Table S1)

**Figure 3:**
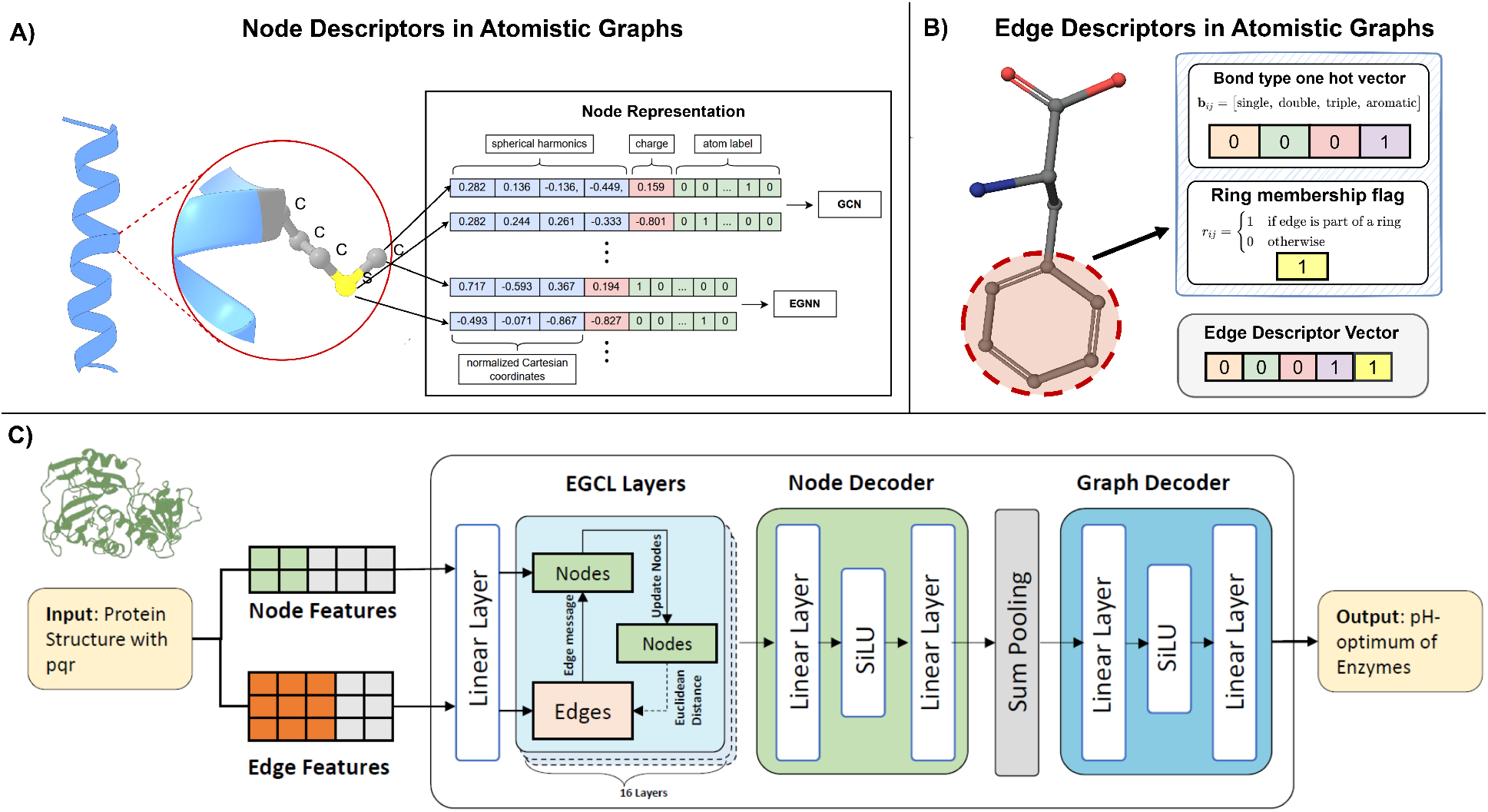
Overview of deep learning framework used for predicting enzyme pH optimum. (**A**) Comparison of node feature vectors constructed for the tested GCNs (top) and EGNNs (bottom). GCNs rely on spherical harmonics, atomic charges, and atom labels, while EGNNs directly incorporate normalized Cartesian coordinates alongside charges and atom labels. (**B**) Each chemical bond is represented by edge features including bond type, ring membership, and the Euclidean distance between atoms. (**C**) Overview of the EGNN architecture used for the pHoptNN model. Node features are first projected into a hidden space through a linear embedding layer. A stack of 16 equivariant graph convolutional layers (EGCLs) iteratively updates node embeddings based on messages that depend on both edge features and inter-atomic distances, while also updating atomic coordinates in an equivariant manner. The resulting node embeddings are refined by a small node-decoder MLP and then aggregated across the graph via sum pooling. A graph-decoder MLP transforms the pooled embedding into the final scalar prediction corresponding to the enzyme’s pH optimum.

To complement this set, we queried UniProt^3^ for all entries containing pH optimum annotations, identifying 5,291 records. After removing duplicate entries that were already present in the BRENDA-derived set and after applying the same quality-control criteria - merging duplicated entries, excluding inconsistent pH ranges, and removing records with unstructured textual pH descriptions - 3,077 unique enzymes were retained. After removing sequences shorter than 32 amino acids or longer than 1,500 amino acids, the combined dataset comprised 12,095 curated enzyme-pH _opt_ records.

For structural annotation, experimentally resolved protein structures were retrieved from the Protein Data Bank (PDB), resulting in 2,822 enzymes with associated coordinates. For enzymes lacking PDB structures, AlphaFold2 models were obtained from the AlphaFold Protein Structure Database, subject to quality thresholds of pTM ≥ 0.5 and median pLDDT ≥ 70; this step contributed 7,650 high-confidence AF2 models. Remaining sequences without PDB or AF2 coverage were modeled using AlphaFold3. AF3 predictions were filtered by removing residues with pLDDT ≤ 70 (up to a maximum of 50% of the protein length) and verifying that template-based AF3 models aligned to their PDB templates with RMSD < 1.0 Å. This procedure yielded 1,216 *de novo* AF3 models and 2,749 AF3 template-refined models.

After these steps, the final EnzyBase12k dataset consisted of 11,615 enzymes with both curated catalytic pH _opt_ values and high-quality three-dimensional structures. The dataset was first divided into training (70%), validation (10%), and test (20%) partitions. The pH opt values were then standardized using the mean and standard deviation computed from the training set only, and the same parameters were applied unchanged to the validation and test sets.

To further assess out-of-distribution generalization, we constructed additional splits based on sequence similarity thresholds of 70%, 50%, and 20%. In parallel, we evaluated functional generalization using an enzyme classification (EC) holdout scheme, in which the model was trained and validated on all EC classes except one and tested exclusively on the held-out class (EC class 4).

### Protein Representation and Feature Engineering

Protein structures from the EnzyBase12k dataset were represented as molecular graphs in which each non-hydrogen atom corresponds to a node and edges denote covalent bonds or spatially proximate interactions. This atomistic graph formulation enables the integration of structural and chemical information into graph neural network (GNN) architectures, where node and edge descriptors form the basis for message passing. Following established practices in molecular graph representation,^34^ we constructed graphs that retain the complete atomic topology of protein structures and encode relevant geometric, electronic, and chemical features.

#### Node Descriptors

Each node *i* ∈ *V* in the graph represents an individual atom from the atomistic topology. To capture both the local chemical identity and the global structural context, we associated each node with a multidimensional feature vector comprising: (i) a one-hot encoding of the atom type **e**_atom_(*i*), (ii) the three Cartesian coordinates **x**_*i*_ ∈ R^3^, and (iii) the atomic partial charge *q*_*i*_ ∈ R, computed with PDB2PQR. ^35^ Optionally, angular information derived from spherical harmonics was included to provide a rotationally informed descriptor of atomic environments. The initial node feature vector is thus defined as:

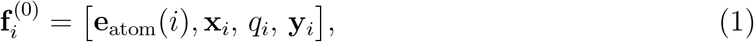

where **y**_*i*_ denotes the set of spherical harmonic features, with normalized coordinates 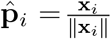.

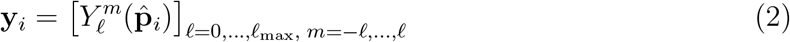

This representation ensures that both local chemical identity and global macromolecular geometry (3D coordinates) are encoded in the graph input.

#### Edge Descriptors

Edges (*i, j*) ∈ *E*encode chemical and geometric relationships between atoms. Covalent bonds were identified using RDKit^36^ parsing rules to ensure correct reconstruction of peptide backbone and amino acid side-chain connectivity. Additional distance-based edges were incorporated to capture non-covalent spatial proximities critical for protein stability and catalysis.

Each edge is described by a five-dimensional feature vector:

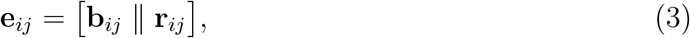

where **b**_*ij*_ ∈ {0, 1}^4^ is a one-hot encoding of the covalent bond type, and **r**_*ij*_ ∈ {0, 1} indicates ring membership. During message passing, geometric information is dynamically incorporated through the squared Euclidean distance between atoms:

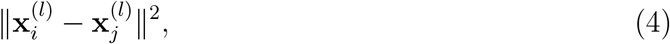

and concatenated with the edge features when forming edge messages,

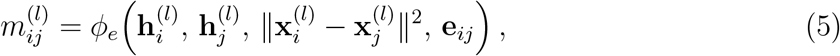

where 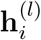 and 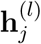 are the current node embeddings at layer *l*, and *ϕ*_*e*_ is a learnable multilayer perceptron. This formulation enables simultaneous integration of covalent connectivity, ring structures, and continuously updated spatial geometry, thereby capturing both local chemical features and long-range structural interactions essential for modeling protein function.

### Model Architecture

The prediction framework employed in this study is based on GNN architectures designed to operate directly on atomistic protein graphs. We evaluated two families of models: (i) conventional graph convolutional networks (GCNs) and (ii) *E*(*n*)-equivariant graph neural networks (EGNNs), which explicitly incorporate geometric constraints by ensuring equivariance to Euclidean symmetries. Below, we summarize the key architectural components and update mechanisms for each model class.

#### Graph Convolutional Networks (GCNs)

As a baseline architecture, we implemented a graph convolutional network following the formulation of Kipf and Welling. ^37^ In this framework, each node’s feature representation is iteratively updated by aggregating features from its local neighborhood. Let 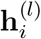 denote the embedding of node *i* at layer *l*. The update rule is:

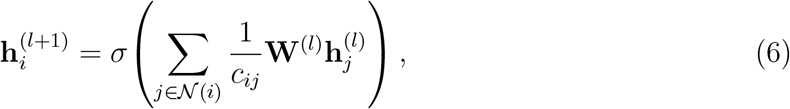

where 𝒩(*i*) is the neighborhood of node 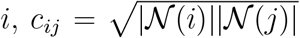 is a normalization factor, **W**^(*l*)^ is a learnable weight matrix, and *σ*(·) is a non-linear activation function.

To introduce geometric sensitivity, node embeddings were augmented with spherical harmonic descriptors (see subsection on Feature Engineering), providing rotationally informative features in otherwise rotation-invariant GCN layers. After *L* convolutional layers, a permutation-invariant readout (mean or sum pooling) aggregates node embeddings into a graph-level representation:

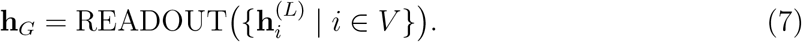

After that, a multi-layer perceptron (MLP) receives the pooled embedding **h**_*G*_ in order to forecast downstream properties, as illustrated in the GCN architecture (Supplementary Figure S1).

#### Equivariant Graph Neural Networks (EGNNs)

To more effectively incorporate protein geometry, we implemented EGNNs,^30^ which ensure that model outputs transform consistently under rotations, translations, and reflections of the input coordinates. EGNNs^30^ circumvent the need for handcrafted geometric features, such as spherical harmonics, and operate directly on Cartesian coordinates. In EGNNs, each node *i* is associated with both a feature embedding 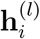 and spatial coordinates 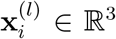, both of which are updated during message passing.

At layer *l*, the message between nodes *i* and *j* is computed as:

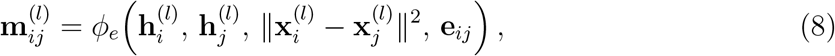

where *ϕ*_*e*_ is a learnable MLP and **e**_*ij*_ denotes the edge descriptor that encodes covalent bond type and ring membership. The squared interatomic distance introduces geometric context while preserving equivariance. The node embeddings are then updated by aggregating incoming messages:

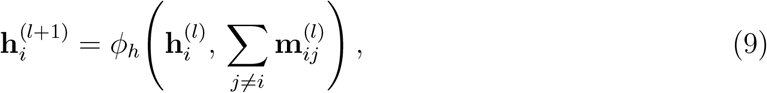

where *ϕ*_*h*_ is another learnable MLP that integrates both local and geometric information. Simultaneously, node coordinates are updated equivariantly as:

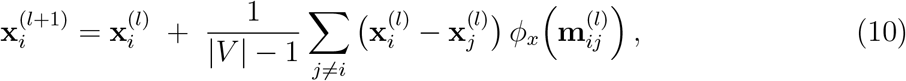

where *ϕ*_*x*_ is a scalar MLP that determines the magnitude of coordinate displacements. This formulation ensures that the model learns geometry-dependent interactions while maintaining *E*(*n*)-equivariance.

After the final EGNN layer, node embeddings are processed through a shallow node-decoder MLP and aggregated using sum pooling to form a graph-level representation:

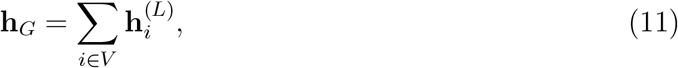

which is subsequently mapped to the enzyme’s predicted pH optimum using a graph-decoder MLP.

In our implementation, the architecture begins with a linear projection layer that maps the input node and edge features to a hidden representation. In *E*(3)-EGNNs, the coordinates are treated separately from the node embeddings, leaving 38 node features in total (37 atom labels and 1 atomic charge). This projection is followed by 16 *E*(3)-equivariant graph convolutional layers (EGCLs). The resulting node embeddings are processed through two linear layers with a SiLU activation function, and then aggregated via sum pooling to obtain a graph-level representation. Finally, a graph decoder consisting of linear layers and a SiLU nonlinearity maps this representation to a single scalar, corresponding to the predicted pH optimum. The best-performing *E*(3)-EGNN employed 16 EGCL layers, the Adam optimizer, a tuned learning rate, and weight decay, with batch size set to 1.

### Evaluation Metrics

Model performance was assessed using a set of complementary regression metrics that together quantify accuracy, error magnitude, and correlation with experimentally measured pH _opt_ values. Specifically, we report the root mean squared error (RMSE), mean squared error (MSE), coefficient of determination (*R*^2^), as well as Pearson’s correlation coefficient (*r*) and Spearman’s rank correlation coefficient (*ρ*).

RMSE and MSE measure the average deviation between predicted (*ŷ*_*i*_) and observed (*y*_*i*_) pH _opt_ values:

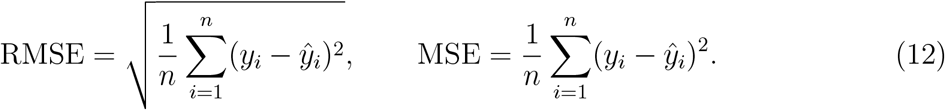

RMSE emphasizes larger errors due to the square-root term, while MSE provides an unsigned measure of overall deviation. The coefficient of determination *R*^2^ quantifies the proportion of variance in the target variable explained by the model:

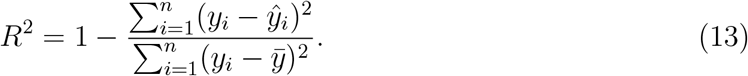

Pearson’s *r* characterizes linear correlation, while Spearman’s *ρ* evaluates the correlation of ranked values, making it less sensitive to outliers and non-linear relationships. During training, MSE and RMSE were monitored to assess convergence, while all five metrics were computed on the held-out test set to provide a comprehensive evaluation of predictive performance.

### Loss Reweighting

The distribution of catalytic pH _opt_ values in EnzyBase12k is highly imbalanced, with most enzymes exhibiting neutral pH optima and substantially fewer examples in strongly acidic or alkaline regions. To mitigate the resulting bias toward densely populated pH bins, we evaluated several loss reweighting strategies (Supplementary Figure S2).

As a straightforward approach we evaluated inverse bin weighting, where the continuous pH range is discretized into fixed-width bins (e.g., 3, 5, or 10 bins) and the weight assigned to each sample is inversely proportional to the number of instances in its bin:

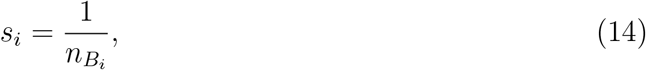

with 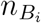 denoting the number of samples in the bin *B*_*i*_ containing *y*_*i*_. Using these weights, a reweighted mean squared error (MSE_reweighted_) is obtained and applied during training and hyperparameter optimization. A moderated variant uses 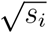 to reduce the sensitivity to very small bins.

In addition, we evaluated distribution smoothing (LDS) as proposed by Yang et al.^38^ LDS convolves the empirical label distribution with a smoothing kernel, producing reweighted density estimates that capture local continuity in the label space while reducing sensitivity to sparse regions. Unlike coarse binning, LDS provides a continuous correction that better accommodates the long-tailed and multimodal characteristics of the pH _opt_ distribution, detailed in Supplementary Figure S2. This approach improves generalization in underrepresented pH ranges while preventing overfitting to rare examples.

### Hyperparameter Optimization

Hyperparameter tuning followed a complementary optimization strategy combining Bayesian optimization and evolutionary search. Initially, Bayesian optimization was used to efficiently explore continuous-valued hyperparameters, including learning rate, weight decay, dropout probability, and hidden dimensionality. This procedure identified high-performing regions of the parameter space and provided well-informed initialization points for a broader search. Subsequently, an evolutionary algorithm, (Supplementary Algorithm S1) was applied to jointly optimize a larger set of 11 hyperparameters, including architectural components (number of EGCL layers, hidden dimension), training parameters (batch size, optimizer settings), and label distribution smoothing parameters (number of bins, kernel size, and smoothing factor). The evolutionary search operated with a population size of 10 per generation and was executed for 1,000 generations. For each candidate configuration, performance was evaluated on the validation set using RMSE as the primary ranking metric. The final hyperparameter configuration was selected based on the lowest validation RMSE and subsequently used to train the pHoptNN model on the full training set. Details of the tested parameter ranges and full optimization trajectories are provided in the see Supplementary Table S2 and S3.

**Table 1.** Final optimized hyperparameter configuration. Complete tested ranges of hyper-parameter can be found in Supplementary Table S3.

| Hyperparameter | Value | Hyperparameter | Value |
| --- | --- | --- | --- |
| *Label distribution smoothing* |  | *EGNN architecture / training* |  |
| number of bins | 65 | number of epochs | 500 |
| kernel size $k_s$ | 1 | learning rate | $6.75 \times 10^{-5}$ |
| $\sigma$ | 0.229 | number of EGCL layers | 16 |
| weighting factor | 0.995 | attention | True |
|  |  | optimizer | Adam |
|  |  | batch size | 1 |
| | | weight decay | $3.1 \times 10^{-7}$ |
|  |  | number of node features | 111 |

### Calculation of Per-Residue Attention Weights

The EGNN employs an edge-level attention mechanism in which each directed atom–atom interaction is assigned a learnable scalar coefficient at every message-passing layer, modulating the strength of information flow during node updates (Supplementary Figure S3). During inference, attention coefficients were extracted from all layers and averaged to obtain a single importance score per edge across the full network depth. Atom-level attention scores were computed by aggregating the averaged attention values of all outgoing edges associated with each atom. These atom-level scores were then mapped to residues using structural annotations, and residue-level attention was defined as the mean attention of all atoms belonging to the same residue.

To facilitate comparison across proteins with different absolute attention scales, attention values were normalized on a per-protein basis prior to downstream analysis. Specifically, atom-level attention scores were divided by the mean attention value of the corresponding protein, resulting in normalized attention values with a protein-wide mean of one. Residue-level attention scores were computed from these normalized atom-level values and used for all subsequent statistical analyses and visualizations. This procedure was used to compare relative attention patterns across proteins and the resulting scores were interpreted as descriptive measures of model emphasis rather than direct evidence of causal residue importance.

For comparative analyses, residue-level attention scores were aggregated across proteins to compute mean attention values for each amino-acid type within discrete enzyme pH bins. In addition, enzymes were grouped into functional classes (proteases, oxidoreductases, and glycosidases), and residue attention was analyzed as a function of structural context by binning residues according to relative solvent accessibility or distance to the annotated active site. Relative solvent accessibility (RSA) for each residue was calculated by determining its solvent-accessible surface area from the enzyme’s 3D structure using FreeSASA ^39^, and dividing this value by maximum solvent accessibility specific to its residue type.

To avoid bias from proteins contributing unequal numbers of residues per bin, attention values were first averaged within each bin on a per-protein basis and subsequently averaged across proteins, with uncertainty estimated from the variation between proteins. This analysis framework allowed systematic assessment of how the model’s learned attention patterns depend on residue identity, pH regime, enzyme class, and spatial proximity to catalytically relevant regions.

## Results and Discussion

### Dataset on Enzyme Structures and pH Optima

To establish a comprehensive foundation for structure-based pH optimum prediction, we compiled the EnzyBase12k dataset, which comprises 11,615 distinct enzymes, spanning all seven major enzyme commission (EC) classes and a broad range of pH optima (Figure 4).

**Figure 4:**
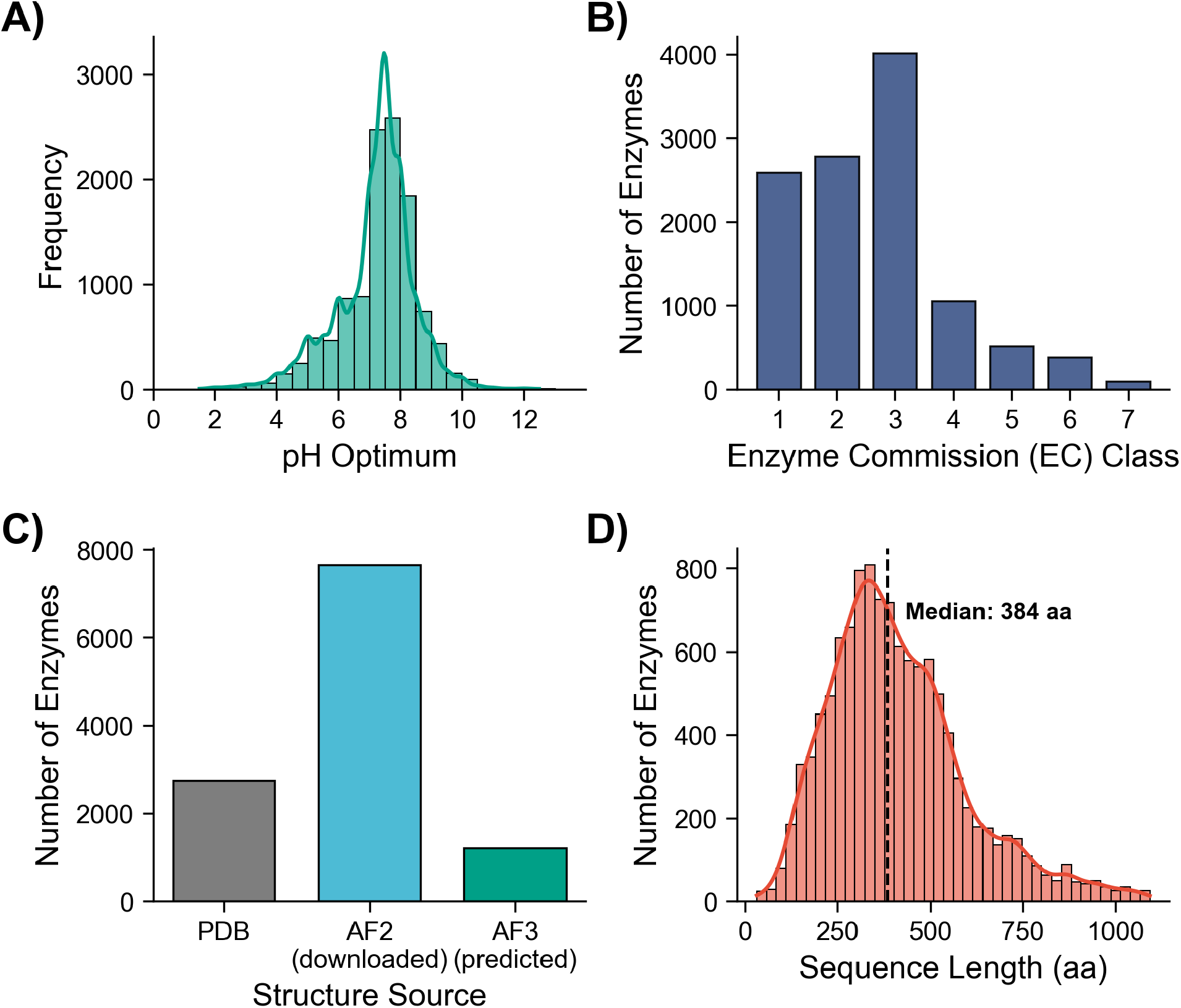
Statistics on the EnzyBase12k dataset. (**A**) Histogram of pH optima. (**B**) Distribution of enzymes in the 7 main EC classes. (**C**) Source of the enzyme structures. (**D**) Distribution of the enzyme sequence length.

As shown in Figure 4A, the distribution of catalytic pH optima is strongly centered around neutral conditions, with a median pH _opt_ of 7.50 and a standard deviation of 1.24 pH units. Only a small subset of enzymes exhibits optimal activity under strongly acidic or alkaline conditions. The overall distribution is slightly skewed toward lower pH values, reflecting the prevalence of enzymes adapted to mildly acidic to neutral environments. A more granular characterization of the pH _opt_ distribution, generated using bin widths of 0.4, 0.3, and 0.2 pH units, is provided in the Supplementary Figure S4. The representation of enzymes across EC classes is similarly heterogeneous (Figure 4B). Hydrolases (EC 3; 4017 enzymes) and transferases (EC 2; 2784 enzymes) constitute the largest fractions of the dataset, whereas ligases (EC 6; 384 enzymes) and translocases (EC 7; 97 enzymes) are comparatively underrepresented.

Figure 4C summarizes the structural provenance of the enzymes included in EnzyBase12k. Of the 11,615 entries, 2,749 enzymes have experimentally determined structures deposited in the Protein Data Bank (PDB). To ensure structural completeness and consistency with UniProt reference sequences, these structures were refined using AlphaFold3 with PDB-derived templates. The remaining 8,866 enzymes lacked experimental structures and were therefore modeled computationally. Of these, 7,650 structures were obtained directly from the AlphaFold Protein Structure Database, while 1,216 enzymes required de novo structure prediction using AlphaFold3 due to the absence of precomputed models.

The sequence length distribution of EnzyBase12k is shown in Figure 4D. The median sequence length is 386 amino acids, with an interquartile range of 285–514 residues, indicating that half of the dataset consists of moderately sized enzymes. The distribution is right-skewed, with proteins longer than 1,200 residues occurring only rarely. Such long sequences are expected to contribute minimally to model training due to their low frequency and limited statistical weight. Dataset composition and structural correlations, including amino acid distributions, species representation, and relationships between sequence length, domain composition, and pH optima, are summarized in the Supporting Information in Figures S5–S6.

### Benchmarking of pHoptNN

Our best-performing equivariant graph neural network, pHoptNN, achieves a mean squared error (MSE) of 0.346 and a root mean squared error (RMSE) of 0.588, together with a strong correlation to experimental pH _opt_ data (Spearman’s *ρ* = 0.78, Pearson’s *r* = 0.89; Table 3). As illustrated in the scatter plot in Figure 5A, predicted pH _opt_ values closely follow the identity line across the entire pH range. Although prediction variance increases modestly under alkaline conditions, residuals remain small overall, underscoring the robustness of the model. The residuals are symmetrically distributed around zero and approximate a Gaussian distribution, with most predictions deviating by less than ±1 pH unit (Figure 5B). This indicates the absence of systematic over- or underestimation. The narrow residual spread and light distribution tails further demonstrate that large prediction errors are rare - an essential property for applications requiring reliable individual predictions.

**Figure 5:**
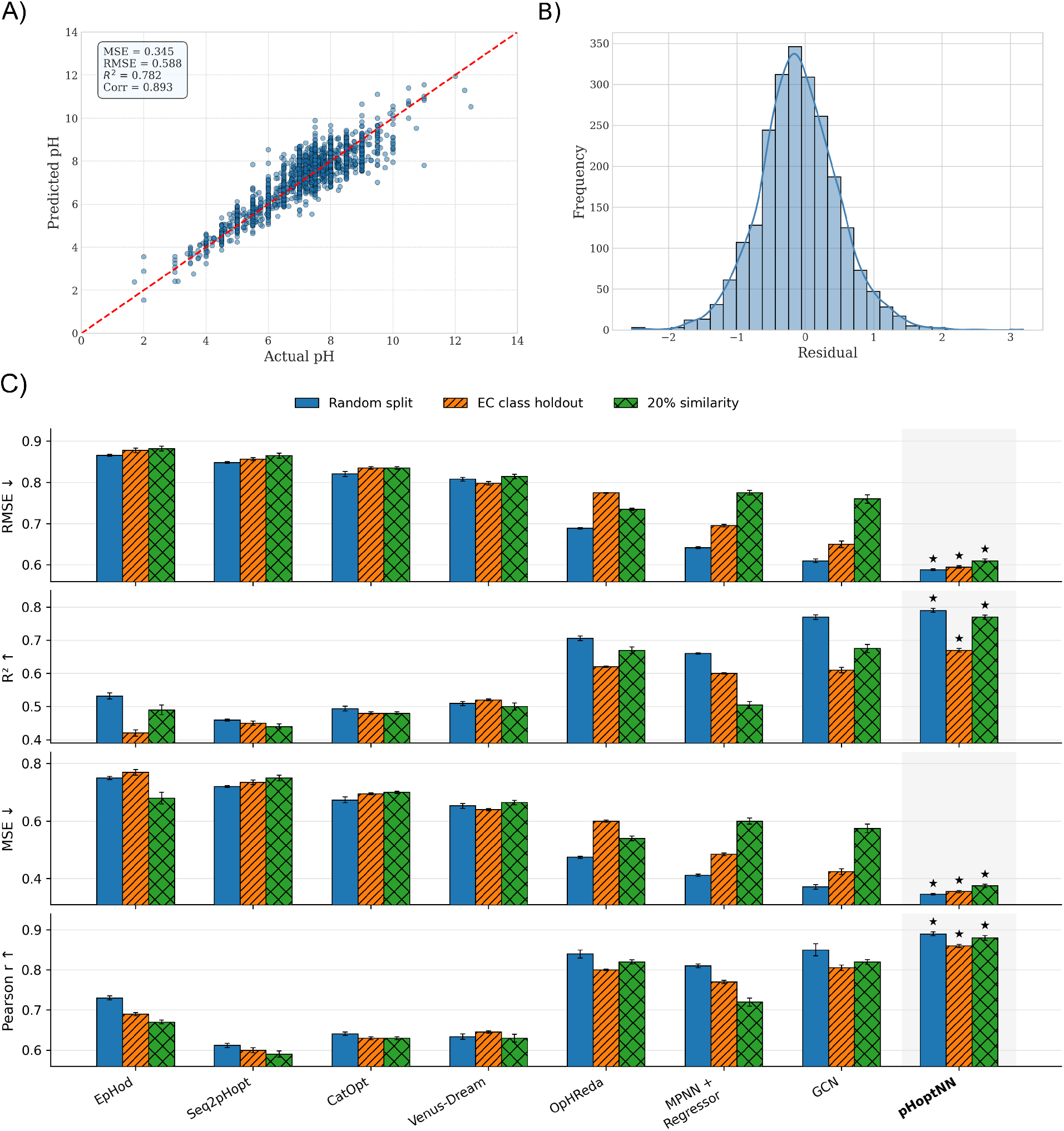
Performance comparison of pHoptNN versus different baseline models. (**A**) Correlation plot for pHoptNN-predicted and actual experimental pH values. (**B**) Residuals of pHoptNN predictions are centered around zero. (**C**) Model performance was assessed using RMSE, *R*^2^, MSE, and Pearson correlation on test sets comprising either EC class 4 enzymes not included in training or enzymes sharing at most 20% sequence similarity with any sequences in the training and validation sets. These results are compared to those obtained under a random data-splitting scheme. Lower RMSE and MSE values indicate better performance, whereas higher *R*^2^ and Pearson correlation values reflect improved predictive accuracy. Bars represent the mean over ten bootstrap iterations, and error bars denote the standard deviation. Detailed numerical results are provided in Tables S5–S7.)

In comparative benchmarking, pHoptNN substantially outperforms the sequence-based baseline EpHod, which attains an RMSE of 0.866, a Spearman’s correlation of 0.63, and a Pearson’s correlation of 0.73. Another sequence-based method, OpHReda, ^14^ improves upon EpHod (RMSE = 0.774, Spearman’s *ρ* = 0.71, Pearson’s *r* = 0.80) but remains inferior to structure-informed models. As structure-based baseline models, we evaluated in-house developed a non-equivariant graph convolutional model based on the Kipf–Welling formulation, augmented with spherical-harmonic geometric descriptors and Set2Set graph-level pooling (Figure S1), as well as hybrid model that uses ProteinMPNN^40^-derived structural embeddings with ESM2 sequence embeddings within a hierarchical regression pipeline (see Supporting method 1.1 for details on architecture and training). The baseline GCN achieved an RMSE of 0.610, outperforming the MPNN+Regressor model, but not surpassing the performance of pHoptNN.

To validate the architectural contribution of the attention mechanism within pHoptNN, we performed an ablation study by removing this module. While the baseline and attention-enable model exhibited comparable aggregate metrics, the attention mechanism proved critical for generalization in extreme regimes, significantly reducing the RMSE in the alkaline range (Supplementary Figure S7). These results demonstrate that explicit incorporation of three-dimensional structural and chemical information provides substantial predictive advantages for enzyme pH optimum estimation.

### Out-of-distribution prediction performance of pHoptNN

To rigorously evaluate the generalization performance of pHoptNN, we conducted benchmark experiments under out-of-distribution conditions that more closely reflect realistic application scenarios. Specifically, we constructed test sets comprising sequences with limited similarity (70%, 50%, or 20%) to any training or validation example. In addition, a functionally distinct subset of enzymes which were excluded from model training and restricted to EC class 4 lyases was used to further challenge model extrapolation.

Figure 5 summarizes the benchmarking results for pHoptNN and the baseline models across three increasingly stringent evaluation settings, with detailed numerical values reported in Supplementary Tables S5–S7. Under the conventional random-split protocol, pHoptNN achieved the best overall performance across all four metrics (Supplementary Table S5). Using a held-out test set composed exclusively of EC class 4 sequences that were entirely excluded from training, pHoptNN maintained strong predictive accuracy and consistently outperformed sequence-based baselines, including Seq2pHopt^11^, CatOpt^12^, Venus-Dream^13^ and OpHReda^14^, as well as the structural and hybrid baseline models (Fig. 5 and Supplementary Table S6).

To further probe generalization to distantly related proteins, we imposed a stringent 20% sequence similarity threshold, thereby minimizing homology between training and test sets. Under these conditions, pHoptNN again demonstrated the strongest overall performance (Fig. 5 and Supplementary Table S7). Notably, the GCN and MPNN+regressor baseline models generally retained higher accuracy than purely sequence-based approaches, such as EpHod^8^ and Seq2pHopt^11^, under low-similarity conditions, underscoring the advantage of incorporating structural information for robust extrapolation.

To systematically examine the dependence of pHoptNN on sequence homology, we evaluated model performance across progressively decreasing similarity thresholds of 70%, 50%, and 20%. As expected, performance declined with decreasing similarity, as reflected by increasing RMSE and MSE and decreasing *R*^2^ and Pearson correlation (Figure 6 and Table S8). This trend is consistent with the increasing evolutionary distance between test and training sequences. Importantly, the decline remained gradual, indicating that pHoptNN retains substantial predictive capacity even under stringent low-similarity conditions. Additional benchmarking results, including analyses across specific pH intervals at the 20% similarity threshold, are provided in Table S9.

**Figure 6:**
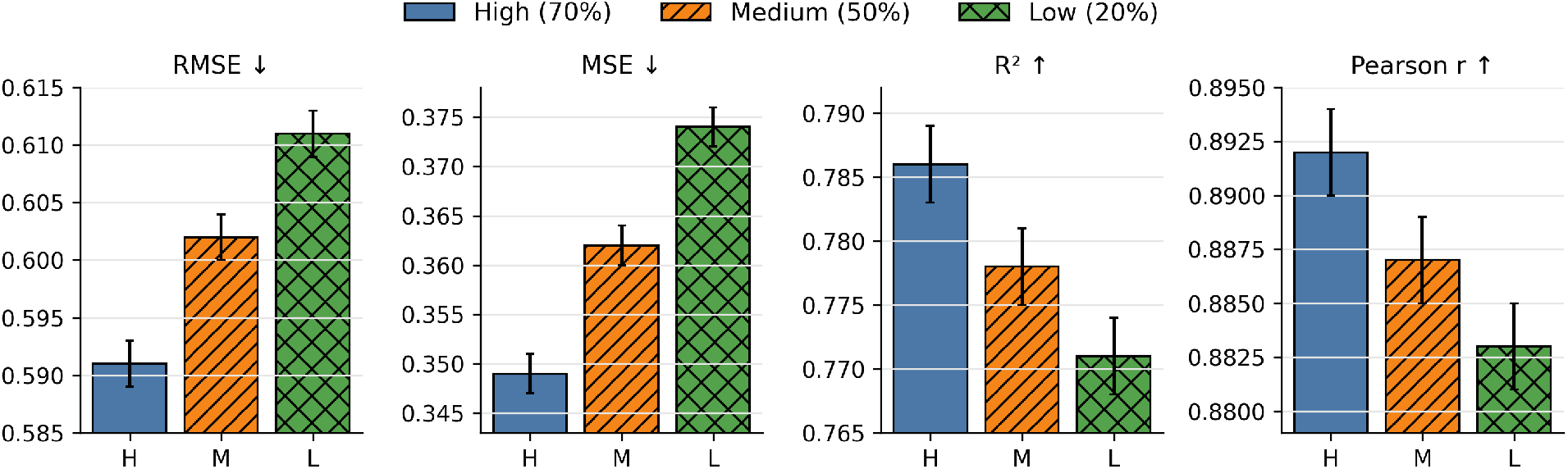
Effect of decreasing sequence similarity between query enzymes and the training data on pHoptNN performance. The model was evaluated on three test sets defined by sequence similarity thresholds of 70% (high), 50% (medium), and 20% (low) relative to any sequence in the training or validation sets. Performance was assessed using RMSE, *R*^2^, MSE, and Pearson correlation. Bars represent the mean over 10 bootstrap iterations, and error bars indicate the standard deviation. Detailed numerical results are provided in Table S8.

To evaluate whether experimentally determined enzyme structures offer a performance advantage over predicted ones, we compared pHoptNN trained on structures from the PDB with an identical model trained on AlphaFold2^32^ structures. Contrary to the expectation that experimental structures might yield superior performance, the model trained on AlphaFold2^32^ structures achieved better overall accuracy than the version trained exclusively on PDB-derived structures (Table S10). This difference is likely attributable, at least in part, to dataset scale: the PDB-based dataset comprised 2,749 experimental structures, whereas the AlphaFold2^32^ dataset included approximately 7,650 structures. The increased size of the AlphaFold2^32^ subset likely enhanced the diversity of structural features available during training, thereby improving model robustness and generalization.

### Stratified Performance Comparison of pHoptNN

Figures 7A and 7B present a stratified comparison of pHoptNN against baseline models across EC classes and pH intervals. Across all EC classes, pHoptNN consistently achieves the lowest prediction error, with particularly pronounced improvements for oxidoreductases (EC 1; RMSE 0.60 for pHoptNN versus 0.91 for EpHod), hydrolases (EC 3; 0.58 versus 0.99), and lyases (EC 4; 0.59 versus 0.84). Even relative to the strongest competing model within each class, pHoptNN maintains performance gains ranging from approximately 3% to 12%. For example, in ligases (EC 6), pHoptNN attains an RMSE of 0.60, compared to 0.68 for the standard GCN, while in translocases (EC 7) the RMSE is reduced from 0.51 to 0.47. The consistency of these improvements across diverse enzyme functions indicates that pHoptNN captures transferable structural determinants of pH dependence rather than overfitting to specific enzyme families.

**Figure 7:**
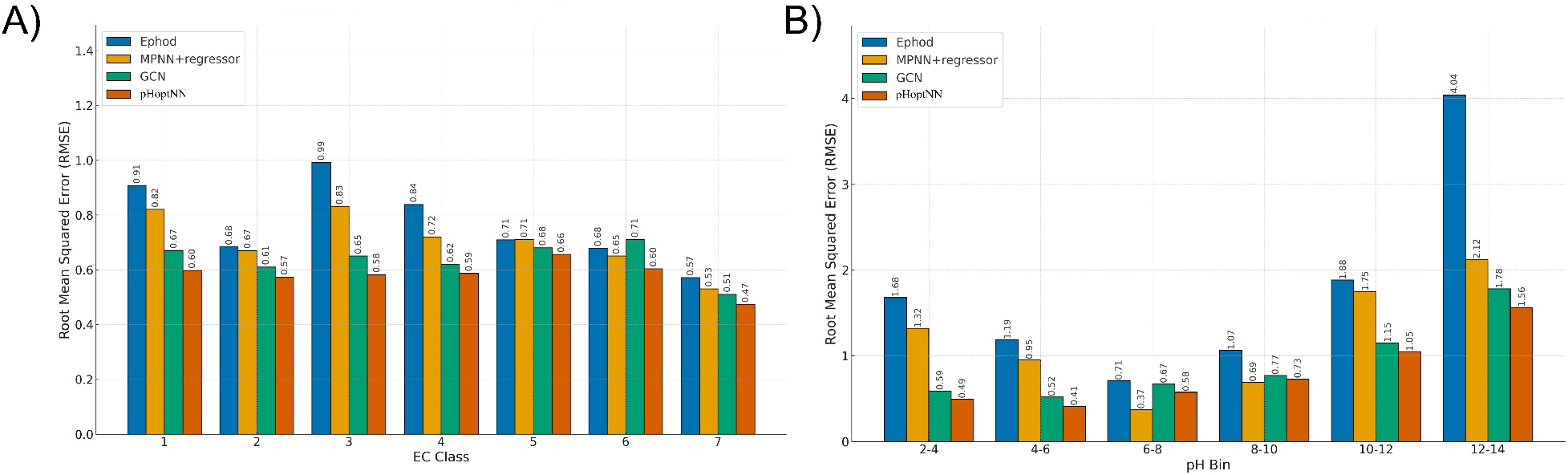
Performance comparison of pHoptNN versus different baseline models. **(A)** RMSE values achieved by pHoptNN (EGNN) and reference methods for each EC class. **(B)** RMSE values achieved by pHoptNN (EGNN) and reference methods for different pH bins.

Performance stratification by pH range further highlights the advantages of the proposed model. pHoptNN exhibits its most substantial gains at the acidic and alkaline extremes of the pH scale, where training data are inherently sparse. In the acidic regime (pH 2–4), the RMSE is reduced from 1.68 for EpHod to 0.49, corresponding to a 71% decrease. Similarly, in the highly alkaline range (pH 12-14), RMSE improves from 4.04 to 1.56, a 61% reduction. In the moderately alkaline range (pH 10-12), pHoptNN lowers the RMSE by 44%, from 1.88 to 1.05. A detailed heatmap illustrating RMSE values, stratified by EC class and pH bin, is provided in Supplementary Figure S9. In contrast, performance differences are less pronounced in the neutral pH range (pH 6-10), where data density is highest. In these bins, the MPNN+regressor model marginally outperforms pHoptNN, suggesting that under conditions of abundant training data, flexible message-passing architectures without explicit equivariant geometric updates can effectively fit dense distributions, while the relative benefit of equivariant reasoning diminishes.

Taken together, the results in Figure 7 demonstrate that pHoptNN delivers substantial and consistent improvements over existing baseline methods. The model achieves strong overall accuracy, robust generalization across enzyme classes, and pronounced gains in underrepresented acidic and alkaline regimes, where conventional approaches perform poorly. Residual analyzes further confirm that the predictions are unbiased and tightly distributed. While performance in the neutral pH range is already high across models, targeted refinements - such as improved loss reweighting strategies - may further reduce the remaining gap to simpler message-passing networks. Overall, pHoptNN emerges as a reliable and broadly applicable framework for enzyme pH optimum prediction.

### Latent-Space Analysis of Representations Learned by pHoptNN

To characterize the internal representations learned by pHoptNN, we analyzed the model’s latent space by visualizing the learned representations using t-SNE,^41^ applied to the final layer node embeddings produced by the trained pHoptNN for each protein, after pooling atom level features into a fixed length graph level embedding with respect to residue identity, chemical class, EC class, and relative solvent accessibility (RSA) (Figure 8). Two-dimensional t-SNE projections reveal that the model organizes residues primarily according to their physicochemical properties, with clear separation of acidic, basic, hydrophobic, and polar amino acids (Figure 8B). In contrast, the t-SNE projection of the EC classes exhibits less distinct clustering (Figure 8C), suggesting that pHoptNN emphasizes local chemical environments and residue-level interactions over global enzyme taxonomy. Nevertheless, the latent representations retain a meaningful structural and functional organization. In particular, the pronounced separation between solvent-exposed and buried residues in the RSA-colored t-SNE projection (Figure 8D) indicates that the model simultaneously encodes global topological features and local chemical context. Together, these findings show that pHoptNN learns chemically interpretable biophysical representations, including electrostatic and topological features (e.g. residue exposure) that are relevant for biocatalysis.

**Figure 8:**
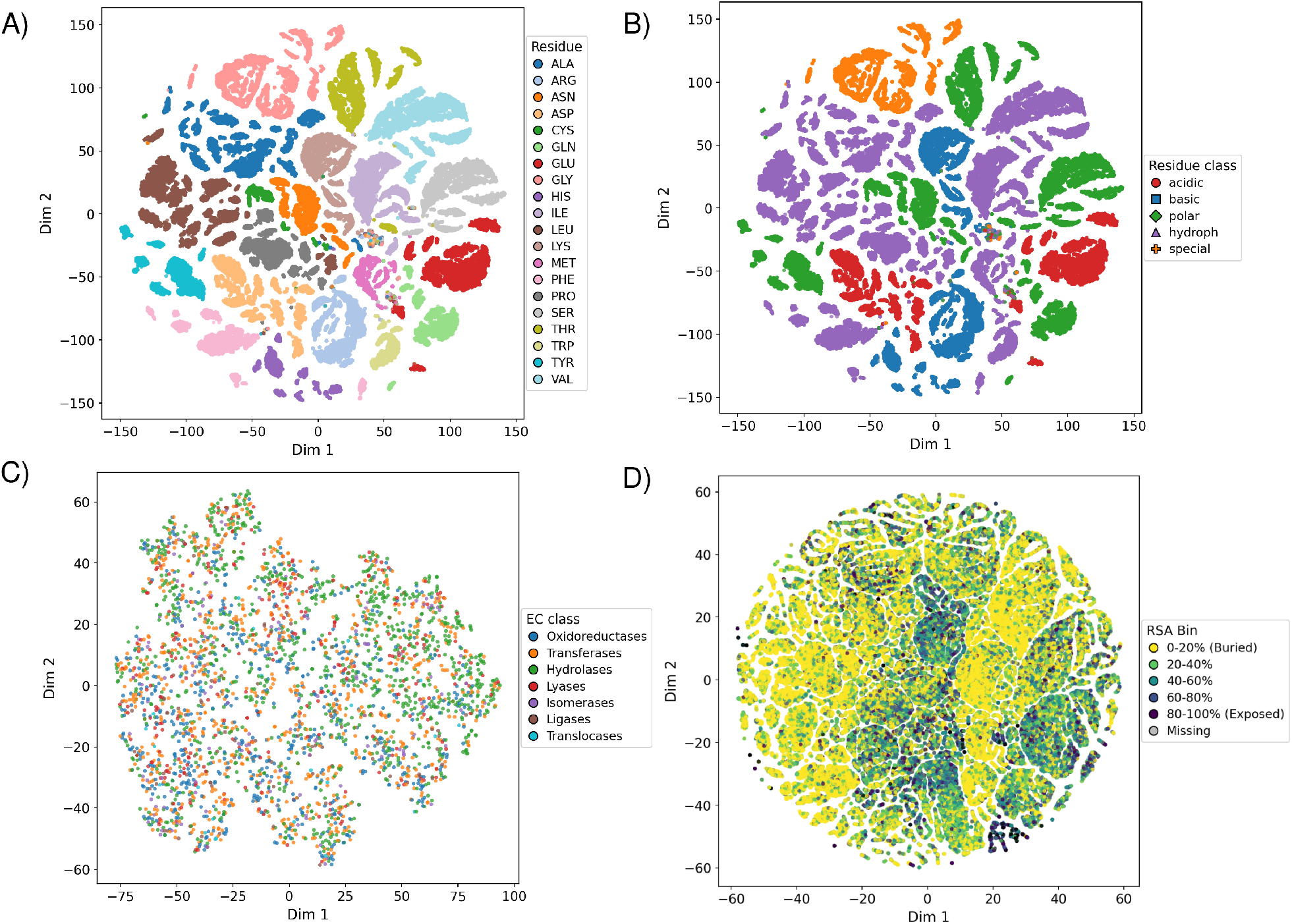
Two-dimensional t-SNE projections of residue- and graph-level embeddings learned by pHoptNN. (**A**) Residue identity clustering reveals grouping of chemically similar amino acids. (**B**) Separation by chemical class reflects learned polarity and charge-based similarities. (**C**) Graph-level embeddings colored by EC class show substantial overlap, indicating that pHoptNN puts less emphasis on global enzyme taxonomy. (**D**) Clustering by relative solvent accessibility (RSA) distinguishes buried from exposed residues, demonstrating that the model encodes both topological depth and solvent exposure.

### pH _opt_-Related Structural Features Learned by pHoptNN

To identify the structural features emphasized by pHoptNN during prediction, attention mechanisms were integrated into the EGCL layers via attention gates (see Supplementary Methods and Figure S3). The resulting residue- and atom-level attention weights can be projected onto corresponding enzyme structures (Figure 9A), enabling direct visualization of regions that contribute most strongly to the pH _opt_ prediction. In these attention maps, residues highlighted in red exhibit high attention—indicating a strong influence on the model’s output—whereas blue regions denote comparatively lower importance.

**Figure 9:**
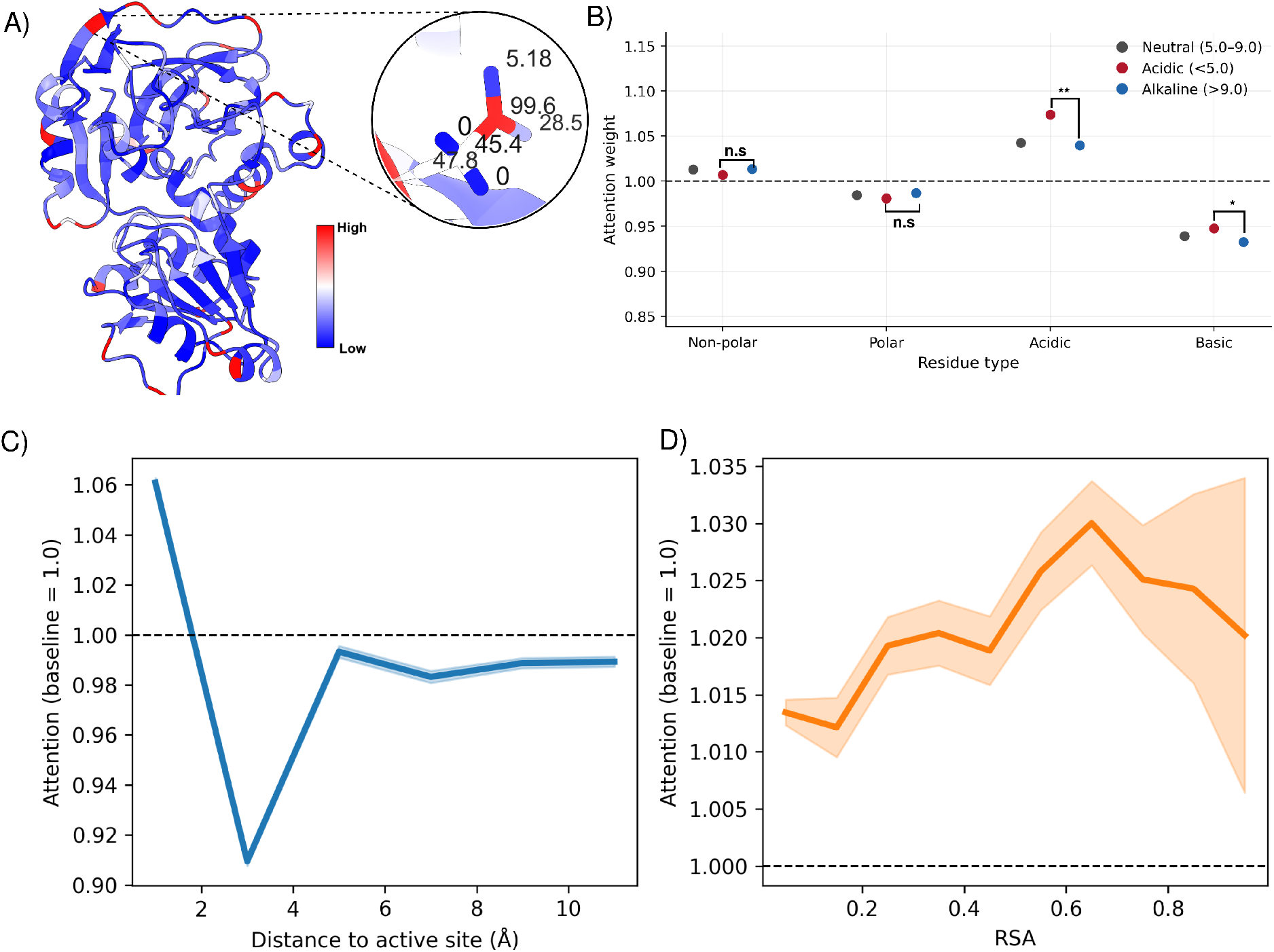
Attention-based analysis reveals physicochemical and structural determinants of enzyme pH optima learned by pHoptNN. (**A**) Residue-wise attention weights projected onto a representative enzyme structure (red, high attention; blue, low attention). The inset highlights atom-level attention scores within an individual residue. (**B**) Mean residue-level attention weights grouped by amino acid chemical class: nonpolar (Val, Leu, Ile, Pro, Phe, Trp, Met, Gly), polar (Ser, Thr, Cys, Tyr, Asn, Gln), acidic (Asp, Glu), and basic (Arg, Lys, His). Two-sided Welch’s t-tests comparing attention weights between acidic and alkaline enzymes reveal statistically significant differences for acidic and basic residues (*P* -values: 4.68 × 10^−45^(**) and 4.695 × 10^−16^(*)), whereas differences for nonpolar and polar residues are not significant (*P* -values: 3.738 × 10^−3^ and 1.204 × 10^−3^). Mean per-residue attention weights for glycosidases (EC 3.2.1.-) plotted as a function of distance to the active site (**C**) and relative solvent accessibility (RSA) (**D**). The dashed line indicates the per-protein normalized mean attention (1.0), while the solid line and shaded region represent the mean attention and ±1 standard deviation calculated across all glycosidase proteins.

Global analysis of residue-level attention across the EnzyBase12k dataset reveals pronounced differences among amino acid types (Figure 9B and Supplementary Figure S10). Ionizable and polar residues, particularly Asp, His, and Asn, receive the highest normalized attention weights, consistent with their established roles in proton binding, hydrogen bonding, and pH buffering within catalytic and surrounding environments. Residues such as Thr and Pro also show elevated attention, reflecting their capacity to participate in hydrogen bonding or nucleophilic interactions that modulate local proton dynamics (Supplementary Figure S10). In contrast, hydrophobic residues—especially Leu, Phe, and Val—tend to receive lower attention, in agreement with their limited involvement in electrostatic or proton-mediated processes. Intriguingly, we found that pHoptNN learned to assign higher attention to negatively charged (acidic) residues in acidic enzymes than alkaline enzymes (Figure 9B). These trends are consistent with biochemical observations that proteins adapted to extreme pH conditions often exhibit an increased abundance of charged residues to maintain stability and catalytic efficiency.

Previous protein engineering studies further support these findings, demonstrating that targeted modification of charged residues, particularly on solvent-exposed surfaces, can systematically shift enzyme pH optima toward more acidic or alkaline values.^42–48^ In addition, residues located near catalytic centers exert a disproportionate influence on the electrostatic environment and protonation states (pKa values) of key catalytic residues, thereby strongly affecting pH-dependent activity.^49–52^

Motivated by these considerations, we analyzed attention weights as a function of relative solvent accessibility (RSA) and distance to the active site (Figure 9C and 9D). On average, pHoptNN assigns higher attention to solvent-exposed residues, consistent with their role in charge stabilization and solvent-mediated proton exchange, as well as to residues in close proximity to the active site. Owing to uncertainties in active-site annotation for many enzymes, the latter analysis was restricted to representative enzyme families with well-defined catalytic residues and sufficient pH _opt_ coverage. Figure 9C illustrates these relationships for glycosidases (EC 3.2.1.-), while extended analyzes for serine (EC 3.4.21.-), cysteine (EC 3.4.22.-), aspartic proteases (EC 3.4.23.-) (Supplementary Figure S11), and oxidoreductases (EC 1.14.-) (Supplementary Figure S11) are provided in the Supplementary Information.

To further assess interpretability, we performed a protease-focused attribution benchmark on 231 enzymes with matched structures and curated catalytic-site annotations. The attention-based strategy preferentially ranked annotated catalytic residues above background, consistent with the high-attention active-site patterns observed in Figure 9c and Supplementary Figure S11. In the supplementary benchmark, this strategy outperformed Integrated Gradients and shuffled-attention controls across AUROC, AUPRC, Hit@1, and top-decile enrichment (Supplementary Figure S12).

We interpret these results as evidence that the model captures residue-level features associated with enzymatic function, rather than as exact catalytic-site prediction. Because attribution scores are not causal explanations and perturbation-based validation remains inconclusive, these findings should be viewed as supportive evidence for biologically meaningful attribution patterns.

### pHoptNN Correctly Predicts Mutation-Induced Shifts in pH _opt_

In addition to analyzing the chemical and structural features learned by pHoptNN, we assessed its ability to support the rational design of enzyme variants with altered pH optima. To this end, we assembled an external benchmark comprising 34 enzyme mutants derived from 18 wild-type enzymes, collected from 14 independent literature sources. ^42–55^ The benchmark includes enzymes from diverse functional classes, including *Aspergillus fumigatus* amine transaminase (EC 2.6.1.18),^53^ human nonpancreatic secretory phospholipase A2 (EC 3.1.1.4),^42^ *Aspergillus niger* GH11 xylanase (EC 3.2.1.8),^49^ *Bacillus sp. YM55-1* aspartase (EC 4.3.1.1),^43^ *Rhodotorula glutinitis* phenylalanine ammonia lyase (EC 4.3.1.24),^50^ *Bacillus subtilis* NADH oxidase (EC 1.6.99.3),^51^ *Bacillus gibsonii* alkaline protease (EC 3.4.21.63),^46^ *Aspergillus oryzae* acidic *β*-glucuronidase (EC 3.2.1.31), ^48^ as well as several xylanases (EC 3.2.1.8) from distinct phylogenetic origins^44,45,47,52,54,55^ (full details are provided in Supplementary Table S5). The reported mutants exhibit pH _opt_ shifts ranging from 0.5 to 2.0 pH units relative to their corresponding wild-type enzymes, and none of the proteins in this benchmark were present in the EnzyBase12k dataset.

pHoptNN correctly predicted the direction of the experimentally observed pH _opt_ shift for 31 out of 34 enzyme mutants. Quantitative agreement within 1.0 pH unit of the experimentally determined pH _opt_ was achieved for 13 of the 18 wild-type enzymes and for 21 of the 34 mutant variants. Moreover, in 15 cases, the pH _opt_ values of both the wild-type enzyme and its corresponding mutant were simultaneously predicted within 1.0 pH unit of the experimental measurements (Supplementary Table S8).

Collectively, these results demonstrate that pHoptNN generalizes beyond wild-type enzymes and is sensitive to subtle physicochemical changes introduced by point mutations. This performance highlights the potential of pHoptNN as a computational tool to assist in the design and screening of enzyme variants with tailored pH optima.

## Conclusion

In this work, we introduced pHoptNN, an *E(n)*-equivariant graph neural network that leverages 3D enzyme structures to predict catalytic pH optima. By representing proteins as atom-level molecular graphs and explicitly encoding geometric, chemical, and electrostatic information, pHoptNN effectively captures the structural determinants underlying pH-dependent enzyme activity. Comprehensive benchmarking on the EnzyBase12k dataset demonstrated that pHoptNN consistently outperforms sequence-based and non-equivariant structure-based baselines, achieving a test-set RMSE of 0.588 pH units. Notably, the model maintains robust performance across enzyme classes and pH regimes, with particularly improvements in acidic and alkaline regions where training data are sparse. At the same time, prediction accuracy at the most extreme pH ranges remains limited by data scarcity and residual label imbalance, highlighting an important opportunity for future improvements as additional experimental data become available.

Beyond predictive accuracy, pHoptNN provides mechanistic insight into enzyme pH adaptation. Attention-based interpretability analyses revealed that the model preferentially focuses on ionizable and polar residues, solvent-exposed regions, and residues proximal to catalytic centers—features that are well established as key contributors to proton transfer, electrostatic stabilization, and pH-dependent catalysis. Furthermore, latent-space analyses showed that pHoptNN learns chemically meaningful representations that reflect residue physicochemical properties and structural context rather than enzyme taxonomy alone.

Looking forward, pHoptNN provides a flexible foundation for further methodological extensions. Incorporation of evolutionary information, explicit residue protonation-state modeling, and refined active-site annotations could further enhance predictive fidelity and biological interpretability. Extending the framework to jointly model additional environmental optima — such as temperature, ionic strength, or solvent composition — would enable a more holistic description of enzyme performance landscapes. Moreover, coupling pHoptNN with generative or optimization-based protein design strategies offers a promising route toward rational enzyme engineering, where predicted pH shifts can directly inform mutation selection and iterative design–build–test cycles.

Overall, pHoptNN illustrates how structure-aware, equivariant deep learning can advance both predictive accuracy and mechanistic understanding in enzymology. By enabling accurate pH optimum prediction and guiding the design of pH-adapted variants, pHoptNN offers a useful computational tool to support enzyme engineering and biocatalysis applications.

## Supporting information

Supplementary informations

## Data and Code Availability

The complete EnzyBase12k dataset used in this study is publicly available at https://doi.org/10.5281/zenodo.15672651. The source code for pHoptNN, including data pre-processing, model training, and interpretability analysis, is freely accessible on GitHub at https://github.com/kuenzelab/pHoptNN. All additional scripts used for benchmarking, visualization, and figure generation are provided within the same repository to ensure full reproducibility of the results.

## Author Contributions

G.K. conceived and supervised the study. C.C. performed the investigation and data curation. R.S. and C.C. developed the model framework, while R.S. carried out the analyses, visualization, and interpretation of results. R.S. prepared the original draft, and both R.S. and G.K. contributed to manuscript review and editing.

## Notes

The authors declare no competing financial interest.

## TOC Graphic

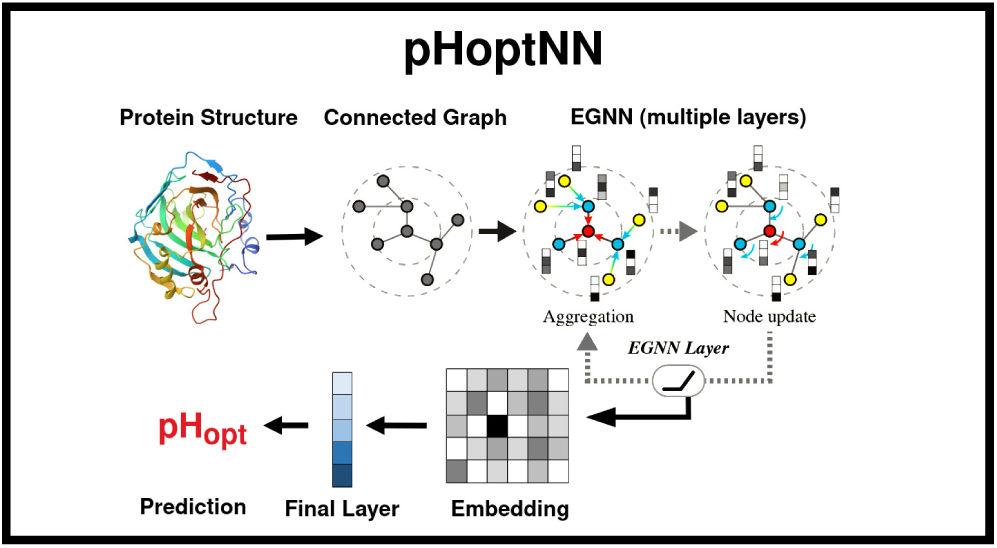

