## Supplementary informations for "Predicting Enzyme pH Optima from Structure Using Equivariant Graph Neural Networks"

### 1. Supplementary Methods

#### 1.1. Model Architecture and Computational Details

All computational experiments were conducted on a high-performance computing (HPC) cluster utilizing NVIDIA H100-SXM5 Tensor Core GPUs (94 GB HBM2e), provided by the Center for Information Services and High Performance Computing (ZIH) at the Technical University Dresden.

The proposed pHoptNN framework employs an $E(n)$-equivariant graph neural network (EGNN) architecture, designed to facilitate equivariant message passing between atomic nodes by integrating both geometric and chemical features. The optimal architecture comprises 16 equivariant graph convolution layers, with a hidden dimensionality of 111 node features per atom. To enhance feature extraction, attention gating was implemented via a

[**]These authors contributed equally to this work.

sigmoid-weighted multi-layer perceptron (MLP) applied to edge messages. Node embeddings were iteratively updated using edge features derived from bond types, ring membership, and interatomic distances. Crucially, coordinate updates were applied at each layer to strictly preserve equivariance.

Training utilized AdamW (with weight decay), Rprop, and SGD optimizers, initialized with default parameters. Hyperparameters, including learning rates, were rigorously tuned via optimization strategies described in Section . To mitigate overfitting, dropout regularization was employed, and weight decay was specifically applied to the AdamW optimizer. Although early stopping criteria were monitored (patience = 50, 100, 150, 200), models were ultimately trained for fixed epochs of 500, 1000, and 5000 to ensure convergence.

The optimal Graph Convolutional Network (GCN) architecture consists of two sequential blocks, each comprising a Graph Convolutional Layer, a Tanh activation function, and a dropout layer, followed by a Set2Set pooling operation. Input nodes are represented by feature vectors of length 42. The first GCN layer projects this to an embedding of size 117, while the second yields an embedding of size 107. The Set2Set global pooling mechanism reduces the $n \times 107$ node feature matrix to a fixed-size graph-level vector of length 214 using a recurrent attention mechanism. Finally, a linear layer maps this vector to a scalar output for regression. This architecture, illustrated in Figure S1, was trained with a learning rate of $2.779 \times 10^{-4}$, a batch size of 1, and an AdamW optimizer with a weight decay of $2.158 \times 10^{-6}$.

For comparative analysis, a hybrid baseline model was constructed, integrating ProteinMPNN[1] with a multi-stage regression pipeline. ProteinMPNN embedded raw protein features into 3D node vectors, with edge information projected into a latent space. These embeddings were augmented with ESM2 embeddings (650M parameters) and physicochemical indices to capture both structural and sequence-based information. This combined representation served as input for an XGBoost regressor. To address data imbalance, Label Distribution Smoothing (LDS) with Distributional Reweighting (DIR) was applied, supplemented by a dynamic sampling strategy to reduce bias toward frequent pH values. The model was validated using K-fold cross-validation. The regression module was organized hierarchically: a gating classifier routed samples to bin-specific XGBoost regressors, while Gaussian Process Regression (GPR) was utilized for extreme tail values. This hierarchical design ensured robust performance across the distribution while providing specialized handling for boundary conditions.

#### 1.2. Attention Implementation in EGNN Layers

Figure S3 illustrates the internal structure of an $E(n)$-equivariant graph convolutional layer (EGCL) equipped with an edge-level attention mechanism. In this architecture, learning is driven by message passing between pairs of atoms rather than by isolated node updates.

At the beginning of the layer, each atom is represented by a feature vector and its Cartesian coordinate. For every neighboring pair of atoms, relative geometric information is computed from their coordinates and combined with node and edge features. This information is processed by an edge MLP to generate a message describing how one atom influences another.

Attention is applied directly to these edge messages. For each directed interaction, the model learns a scalar gating value that modulates the strength of the corresponding message. This allows the network to emphasize chemically or structurally relevant interactions while down-weighting less informative ones. Importantly, attention operates on interactions rather than atoms, enabling the model to distinguish which specific neighbors contribute most strongly to a given update.

The attention-weighted messages are aggregated at the receiving atom and used to update node features, capturing the influence of the local interaction environment. In parallel, the same modulated messages contribute to coordinate updates. Because coordinate updates depend only on relative atomic positions, the model remains equivariant to global translations and rotations.

Defining attention at the edge level is a natural choice for EGNNs, as molecular and protein properties emerge from pairwise interactions such as bonding, electrostatics, and spatial proximity. Applying attention at the node level would collapse all neighboring effects into a single scalar and obscure which interactions are most influential. Edge-level attention preserves this relational structure while still allowing post hoc aggregation for interpretation.

For interpretability, edge-level attention values can be aggregated after inference to obtain atom- or residue-level importance scores. In this work, attention coefficients were averaged across layers and summed over edges connected to each atom, and residue-level scores were obtained by averaging atom-level values. These scores were subsequently mapped to structural representations for visualization.

#### 1.3. Loss Reweighting Analysis

Figure S2 depicts the impact of reweighting schemes on the target distribution using Kernel Density Estimation (KDE). The black curve represents the raw $pH_{\mathrm{opt}}$ distribution. The orange and red curves illustrate inverse binning with three bins, which yields only partial smoothing. In contrast, increasing the bin count (e.g., to 10) results in a significantly flatter effective distribution. This reduction in bias towards overrepresented pH ranges is critical for facilitating balanced training dynamics.

#### 1.4. Hyperparameter Optimization Strategies

EGNN optimization was achieved through two complementary strategies: an evolutionary algorithm (EA) for broad exploration and Bayesian optimization for focused refinement.

To optimize the hyperparameters of the graph neural networks used in this thesis, an evolutionary algorithm (EA) was implemented. Inspired by the principles of biological evolution, the algorithm searches for high-performing configurations through selection, recombination, and mutation. Over many generations, the evolutionary algorithm gradually improves initially suboptimal solutions. EAs are applicable to a broad range of optimization problems. In this thesis, one objective was to find a GNN architecture, i.e. a hyperparameter configuration, that minimizes the model's prediction error on the test dataset. The evolutionary algorithm was run with a population size of 10 individuals per generation, where each individual represented exactly one hyperparameter configuration, and for 1,000 generations. The performance of an individual was evaluated based on the inverse test loss, that is lower MSE losses correspond to higher fitness. Initially, a population of 10 individuals was created randomly. In each generation, four out of the 10 individuals from the previous generation were selected as parents using the roulette wheel method. The selection probability of each individual was set by its fitness divided by the total fitness of the population where fitness was defined as the inverse of the MSE loss. Thus, better-performing individuals were more likely to be selected. This selection strategy guided the search through the hyperparameter space to balance exploration and exploitation. At the transition to the next generation, the fittest individual was preserved unchanged (elitism with count one), while the remaining nine individuals were created through crossover and mutation. During crossover, each hyperparameter was randomly inherited from one of the four selected parent configurations.

After crossover, 1 to 12 hyperparameters of each individual were randomly selected for mutation. With a 10% probability, an extreme mutation was applied, allowing larger changes to encourage exploration of the hyperparameter space. Hyperparameters allowed to mutate included learning rate, number of epochs, dropout, optimizer, batch size, hidden layer sizes, pooling method, kernel size, Gaussian width, and the number of histogram bins. Some mutations were dependent on others, e.g., weight decay was scaled with respect to the learning rate.

The procedure is formalized in Algorithm .

**Algorithm S1:** Evolutionary Hyperparameter Optimization

**Input:** Initial population $P$, number of generations $G$

**Output:** Optimal hyperparameter configuration $p^*$

1. Initialize population $P$
2. For each individual $p \in P$, evaluate fitness $f(p)$
3. For $t = 1$ to $G$:
4. Select parents $M \subset P$ via roulette-wheel selection
5. Generate offspring $P'$ via crossover and mutation of $M$
6. Evaluate fitness $f(p)$ for all $p \in P'$
7. Apply elitism: retain best $p \in P$
8. Update population $P \leftarrow P'$
9. End For
10. **Return** best-performing individual $p^*$

To complement the EA, Bayesian optimization was conducted using the Tree-structured Parzen Estimator (TPE) framework. This approach adaptively models the objective function to balance exploration and exploitation. The search space encompassed architectural and training parameters, including epoch count (200–1000), optimizer choice (Adam, AdamW, RMSprop), log-uniform learning rate ($10^{-7}$–$10^{-2}$), and weight decay ($10^{-6}$–$10^{-3}$). Structural parameters included layer count (2–20), hidden features (64–384), and attention mechanisms. Kernel parameters ($k_s$, $\sigma$) and weighting factors were also tuned. Five-fold cross-validation loss served as the objective function, allowing the Bayesian search to efficiently converge toward stable, high-performance configurations.

This dual optimization strategy ensured both the diversity of candidate models and the precise tuning of promising architectures, resulting in robust generalization and minimized validation error across all folds.

### 2. Supplementary Tables

**Table S1.** Dataset reduction and filtering statistics at each curation step.

| **Step** | **Entries** |
|---|---|
| 1.1 BRENDA database download | 96,955 |
| 1.2 Filtering for pH optimum and valid UniProt IDs | 11,001 |
| 1.3 Merging of UniProt ID duplicates | 10,296 |
| 1.4 Validation of UniProt ID accessibility | 9,766 |
| 1.5 Averaging pH (range $\leq$ 1); discarding high-variance entries | 9,537 |
| 2.1 Acquisition of UniProt pH optimum data | 5,291 |
| 2.2 Averaging pH (range $\leq$ 1); discarding high-variance entries | 4,518 |
| 2.3 Removal of UniProt IDs present in BRENDA | 3,256 |
| 2.4 Removal of entries with string processing errors | 3,077 |
| 3. Sequence length truncation (32–1500 amino acids) | 12,095 |
| 4. AlphaFold (AF) download and AF3 calculations | 9,261 + 2,805* |
| 5. Final filtering of AF models | 8,866 + 2,749* |
| **Total Valid Entries** | **11,615** |

* Denotes entries with associated PDB structural information.

**Table S2.** Top hyperparameter configurations identified via evolutionary and Bayesian optimization.

| **Epochs** | **LR** | **Attn** | **Layers** | **Opt** | **BS** | **Loss** | **WD** | **Dim** | $k_s$ | $\sigma$ | **Wt** | **Bins** |
|---|---|---|---|---|---|---|---|---|---|---|---|---|
| 1000 | $1.83 \times 10^{-5}$ | F | 10 | Adam | 1 | LDS | $1.85 \times 10^{-6}$ | 96 | 8 | 1.27 | 0.17 | 51 |
| 1000 | $1.43 \times 10^{-5}$ | F | 6 | AdamW | 1 | LDS | $2.08 \times 10^{-6}$ | 64 | 9 | 1.07 | 0.46 | 111 |
| 1000 | $1.42 \times 10^{-5}$ | F | 6 | AdamW | 1 | LDS | $1.83 \times 10^{-6}$ | 64 | 9 | 1.11 | 0.50 | 124 |
| 1000 | $1.25 \times 10^{-5}$ | T | 12 | AdamW | 1 | LDS | $9.43 \times 10^{-5}$ | 384 | 15 | 1.25 | 0.28 | 28 |
| 1000 | $5.96 \times 10^{-6}$ | T | 8 | AdamW | 1 | LDS | $1.20 \times 10^{-4}$ | 288 | 19 | 0.70 | 0.28 | 112 |
| 1000 | $4.43 \times 10^{-5}$ | T | 18 | Adam | 1 | LDS | $2.00 \times 10^{-7}$ | 103 | 1 | 1.24 | 0.73 | 115 |
| 1000 | $6.75 \times 10^{-5}$ | T | 16 | Adam | 1 | LDS | $3.10 \times 10^{-7}$ | 111 | 1 | 0.23 | 1.00 | 65 |
| 1000 | $5.60 \times 10^{-5}$ | F | 17 | Adam | 1 | LDS | $1.20 \times 10^{-7}$ | 119 | 9 | 1.49 | 0.15 | 91 |
| 1000 | $5.41 \times 10^{-5}$ | F | 15 | RMS | 1 | LDS | $2.10 \times 10^{-7}$ | 81 | 9 | 0.16 | 0.38 | 61 |
| 1000 | $1.11 \times 10^{-4}$ | F | 19 | RMS | 1 | LDS | $2.10 \times 10^{-7}$ | 115 | 5 | 0.31 | 0.42 | 60 |

*Note:* Attn = Attention; Opt = Optimizer; BS = Batch size; WD = Weight decay; Dim = Hidden node features; Wt = Weighting factor; LDS = Label Distribution Smoothing.

**Table S3.** Hyperparameter search space configuration. Ranges denote min–max values for uniform sampling, and lists denote categorical choices.

| Category | Hyperparameter | Search Space / Values |
|---|---|---|
| ***Optimization*** | | |
| | Optimizer | {Adam, AdamW, RAdam, RMSprop, SGD, Adamax} |
| | Learning Rate (LR) | $10^{-7}$ – $10^{-1}$ |
| | Weight Decay (WD) | $10^{-7}$ – $10^{-2}$ |
| | Batch Size (BS) | {1, 8, 16, 32, 64} |
| | Epochs | 100 – 1000 |
| ***Model Architecture*** | | |
| | Layers | 2 – 16 |
| | Hidden Dimension | 32 – 128 |
| | Attention | {True, False} |
| | Kernel Size ($k_s$) | 3 – 15 |
| ***Loss Function (LDS)*** | | |
| | Loss Variant | {LDS, $\text{LDS}_{\text{extreme}}$, None} |
| | Sigma ($\sigma$) | 0.1 – 2.0 |
| | Weight (Wt) | 0.1 – 2.0 |
| | Bins | 1 – 150 |

**Table S4.** MSE for predicted pH optima on EpHod test dataset vs. reduced EnzyBase12k dataset (excluding EpHod training data).

| | EpHod Test Dataset | | EnzyBase12k (Reduced) | |
|---|---|---|---|---|
| **pH Bin** | **MSE** | **# Samples** | **MSE** | **# Samples** |
| 0–2 | 6.878 | 1 | 7.763 | 2 |
| 2–4 | 3.071 | 13 | 4.923 | 76 |
| 4–6 | 1.406 | 180 | 1.965 | 724 |
| 6–8 | 0.548 | 963 | 0.637 | 3,138 |
| 8–10 | 0.785 | 445 | 1.065 | 1,672 |
| 10–12 | 3.114 | 21 | 4.968 | 85 |
| **Total** | **0.773** | **1,623** | **1.068** | **5,697** |

*Note: Values represent mean ± standard deviation estimated via 10 bootstrap iterations.*

**Table S5.** Overall performance metrics for pHoptNN and benchmark methods across the full pH range.

| Model | RMSE ↓ | MSE ↓ | $R^2$ ↑ | Pearson's $r$ ↑ |
|---|---|---|---|---|
| EpHod[2] | $0.866 \pm 0.002$ | $0.750 \pm 0.005$ | $0.532 \pm 0.009$ | $0.730 \pm 0.005$ |
| Seq2pHopt[3] | $0.849 \pm 0.002$ | $0.721 \pm 0.003$ | $0.459 \pm 0.003$ | $0.612 \pm 0.005$ |
| CatOpt[4] | $0.821 \pm 0.006$ | $0.674 \pm 0.010$ | $0.494 \pm 0.007$ | $0.641 \pm 0.004$ |
| Venus-Dream[5] | $0.808 \pm 0.005$ | $0.653 \pm 0.008$ | $0.509 \pm 0.006$ | $0.634 \pm 0.007$ |
| OpHReda[6] | $0.689 \pm 0.002$ | $0.475 \pm 0.004$ | $0.706 \pm 0.007$ | $0.840 \pm 0.010$ |
| MPNN + Regressor | $0.642 \pm 0.002$ | $0.412 \pm 0.004$ | $0.660 \pm 0.002$ | $0.810 \pm 0.005$ |
| GCN | $0.610 \pm 0.004$ | $0.372 \pm 0.008$ | $0.790 \pm 0.007$ | $0.890 \pm 0.015$ |
| **pHoptNN** | $\mathbf{0.588 \pm 0.002}$ | $\mathbf{0.346 \pm 0.003}$ | $\mathbf{0.790 \pm 0.006}$ | $\mathbf{0.890 \pm 0.005}$ |

*Note: Values represent mean ± standard deviation estimated via 10 bootstrap iterations.*

**Table S6.** Benchmarking results on the held-out EC class 4 dataset for out-of-distribution evaluation. Best performances are highlighted in **bold**.

| Model | RMSE ↓ | MSE ↓ | $R^2$ ↑ | Pearson's $r$ ↑ |
|---|---|---|---|---|
| EpHod[2] | $0.878 \pm 0.007$ | $0.771 \pm 0.012$ | $0.422 \pm 0.009$ | $0.689 \pm 0.005$ |
| Seq2pHopt[3] | $0.855 \pm 0.004$ | $0.731 \pm 0.007$ | $0.451 \pm 0.005$ | $0.601 \pm 0.005$ |
| CatOpt[4] | $0.834 \pm 0.002$ | $0.696 \pm 0.003$ | $0.478 \pm 0.003$ | $0.630 \pm 0.004$ |
| Venus-Dream[5] | $0.798 \pm 0.002$ | $0.637 \pm 0.003$ | $0.521 \pm 0.002$ | $0.642 \pm 0.002$ |
| OpHReda[6] | $0.775 \pm 0.001$ | $0.601 \pm 0.002$ | $0.619 \pm 0.001$ | $0.800 \pm 0.001$ |
| MPNN + Regressor | $0.695 \pm 0.002$ | $0.483 \pm 0.003$ | $0.601 \pm 0.002$ | $0.773 \pm 0.003$ |
| GCN | $0.651 \pm 0.009$ | $0.424 \pm 0.012$ | $0.611 \pm 0.007$ | $0.803 \pm 0.006$ |
| **pHoptNN** | $\mathbf{0.594 \pm 0.005}$ | $\mathbf{0.352 \pm 0.006}$ | $\mathbf{0.672 \pm 0.006}$ | $\mathbf{0.859 \pm 0.005}$ |

*Note: Values represent mean ± standard deviation estimated via 10 bootstrap iterations.*

**Table S7.** Benchmarking results on the 20% sequence similarity cluster to assess generalization to distantly related sequences.

| Model | RMSE ↓ | MSE ↓ | $R^2$ ↑ | Pearson's $r$ ↑ |
|---|---|---|---|---|
| EpHod[2] | $0.883 \pm 0.005$ | $0.679 \pm 0.020$ | $0.491 \pm 0.015$ | $0.670 \pm 0.003$ |
| Seq2pHopt[3] | $0.865 \pm 0.005$ | $0.748 \pm 0.009$ | $0.439 \pm 0.006$ | $0.594 \pm 0.006$ |
| CatOpt[4] | $0.835 \pm 0.002$ | $0.697 \pm 0.003$ | $0.477 \pm 0.003$ | $0.630 \pm 0.003$ |
| Venus-Dream[5] | $0.815 \pm 0.003$ | $0.664 \pm 0.005$ | $0.501 \pm 0.010$ | $0.629 \pm 0.007$ |
| OpHReda[6] | $0.735 \pm 0.002$ | $0.540 \pm 0.004$ | $0.666 \pm 0.010$ | $0.821 \pm 0.003$ |
| MPNN + Regressor | $0.773 \pm 0.003$ | $0.598 \pm 0.005$ | $0.506 \pm 0.005$ | $0.719 \pm 0.007$ |
| GCN | $0.759 \pm 0.010$ | $0.576 \pm 0.015$ | $0.675 \pm 0.009$ | $0.822 \pm 0.005$ |
| **pHoptNN** | $\mathbf{0.611 \pm 0.002}$ | $\mathbf{0.374 \pm 0.002}$ | $\mathbf{0.771 \pm 0.003}$ | $\mathbf{0.883 \pm 0.002}$ |

*Note: Values represent mean ± standard deviation estimated via 10 bootstrap iterations.*

**Table S8.** Benchmarking results of pHoptNN across decreasing sequence identity thresholds.

| **Similarity** | **Identity** | **RMSE ↓** | **MSE ↓** | $R^2$ ↑ | **Pearson's** $r$ ↑ |
|---|---|---|---|---|---|
| High | 70% | $0.591 \pm 0.002$ | $0.349 \pm 0.002$ | $0.786 \pm 0.003$ | $0.892 \pm 0.002$ |
| Medium | 50% | $0.602 \pm 0.002$ | $0.362 \pm 0.002$ | $0.778 \pm 0.003$ | $0.887 \pm 0.002$ |
| Low | 20% | $\mathbf{0.611 \pm 0.002}$ | $\mathbf{0.374 \pm 0.002}$ | $\mathbf{0.771 \pm 0.003}$ | $\mathbf{0.883 \pm 0.002}$ |

*Note: Values represent mean ± standard deviation estimated via 10 bootstrap iterations.*

**Table S9.** Benchmarking results of pHoptNN overall and across specific pH intervals for the Low (20%) sequence identity threshold.

| **Data Subset** | **RMSE ↓** | **MAE ↓** | **Pearson's** $r$ ↑ | **Spearman's** $\rho$ ↑ |
|---|---|---|---|---|
| **Overall** | $\mathbf{0.611 \pm 0.012}$ | $\mathbf{0.468 \pm 0.008}$ | $\mathbf{0.884 \pm 0.006}$ | $\mathbf{0.806 \pm 0.010}$ |
| pH 2–4 | $0.623 \pm 0.090$ | $0.465 \pm 0.064$ | $0.550 \pm 0.072$ | $0.601 \pm 0.098$ |
| pH 4–6 | $0.465 \pm 0.018$ | $0.353 \pm 0.012$ | $0.764 \pm 0.029$ | $0.729 \pm 0.038$ |
| pH 6–8 | $0.576 \pm 0.010$ | $0.450 \pm 0.007$ | $0.725 \pm 0.011$ | $0.641 \pm 0.019$ |
| pH 8–10 | $0.724 \pm 0.023$ | $0.559 \pm 0.015$ | $0.405 \pm 0.042$ | $0.373 \pm 0.051$ |
| pH 10–12 | $0.961 \pm 0.133$ | $0.731 \pm 0.108$ | $0.749 \pm 0.064$ | $0.731 \pm 0.061$ |

*Note: Values represent mean ± standard deviation estimated via 10 bootstrap iterations.*

**Table S10.** Comparative analysis of pHoptNN performance using experimental (PDB) vs. predicted (AlphaFold2) structural data.

| **Structure Type** | **Source** | **RMSE ↓** | **MSE ↓** | $R^2$ ↑ | **Pearson's** $r$ ↑ |
|---|---|---|---|---|---|
| Experimental | PDB (Crystal) | $0.732 \pm 0.002$ | $0.536 \pm 0.003$ | $0.607 \pm 0.004$ | $0.785 \pm 0.003$ |
| Predicted | AlphaFold2 | $\mathbf{0.613 \pm 0.005}$ | $\mathbf{0.376 \pm 0.006}$ | $\mathbf{0.721 \pm 0.007}$ | $\mathbf{0.849 \pm 0.005}$ |

*Note: Values represent mean ± standard deviation estimated via 10 bootstrap iterations.*

**Table S11.** Benchmarking of pH $_{opt}$-shift predictions by pHoptNN for experimentally characterized enzyme variants. The table reports the enzyme name, source organism, UniProt and PDB identifiers, introduced mutations, experimentally determined pH $_{opt}$ values for the wild-type (WT) and mutant (Mut) enzymes, and the corresponding pH $_{opt}$ shift (WT → Mut) predicted by pHoptNN.

| Ref. | Enzyme | Organism | UniProt / NCBI | PDB | Mutations | WT pH | Mut pH | pHopt Shift |
|---|---|---|---|---|---|---|---|---|
| 7 | Amine transaminase ATA-Afu | *Aspergillus fumigatus* | Q4WH08 | 4CHI | E49Q | 8.5 | 7.5 | 8.21 → 7.2 |
| 7 | Amine transaminase ATA-Ate | *Aspergillus terreus* | Q0C8G1 | – | Q51E | 7 | 8 | 7.61 → 7.8 |
| 7 | Amine transaminase ATA-Gze | *Gibberella zeae* | I1RDQ2 | 4CE5 | Q49E | 7.5 | 8.5 | 7.24 → 7.35 |
| 8 | non-pancreatic secretory phospholipase A2 hnpsPLA2 | *Homo sapiens* | P14555 | 1KQU | R100E | 8.5 | 7.2 | 7.95 → 7.1 |
| 9 | GH11 xylanase AnXynB | *Aspergillus niger* | P55330 | – | Q178R | 5.5 | 6 | 6.32 → 6.32 |
| 10 | Aspartase AspBm | *Bacillus sp. YM55-1* | Q9LCC6 | 3R6V | K19E, N87E, N125D, S133D, Q262E, N451E | 9 | 8 | 8.32 → 8.2 |
| 11 | Endo-1,4-beta-xylanase | *Bacillus circulans* | P09850 | 1XNB | Y5R | 6.5 | 6 | 6.7 → 6.6 |
| 11 | Endo-1,4-beta-xylanase | *Bacillus circulans* | P09850 | 1XNB | V37R | 6.5 | 5 | 6.7 → 5.69 |
| 11 | Endo-1,4-beta-xylanase | *Bacillus circulans* | P09850 | 1XNB | N63R | 6.5 | 6 | 6.68 → 6.68 |
| 12 | Phenylalanine ammonia-lyase RgPAL | *Rhodotorula glutinis JN-1* | AUQ35650.1 | – | Q137E | 9 | 8 | 8.32 → 7.9 |
| 13 | NADH oxidase BsNox | *Bacillus subtilis* | P81102 | – | N20D, N116E | 9 | 7 | 7.27 → 7.18 |
| 14 | Alkaline xylanase Xyn11A-LC | *Bacillus sp. SN5* | AGL92443.1 | 4IXL | E135V | 7.5 | 8 | 8.38 → 8.9 |
| 14 | Alkaline xylanase Xyn11A-LC | *Bacillus sp. SN5* | AGL92443.1 | 4IXL | E135R | 7.5 | 8.5 | 8.38 → 8.92 |
| 15 | GH11 xylanase | *Caldicellulosiruptor bescii* | WP_015906728.1 | – | S56D, A166E, D176Y, Q177E | 6.5 | 5 | 5.77 → 5.73 |
| 16 | GH11 xylanase BsXynA | *Bacillus subtilis* | P18429 | 1XXN, 2DCY | Q7H, G13R, S22P, S31Y, T44A, I51V, I107L, S179C | 6 | 6.5 | 5.5 → 5.95 |
| 16 | GH11 xylanase AnXynB | *Aspergillus niger* | P55330 | – | D117N | 5 | 5.5 | 7.56 → 7.65 |
| 16 | GH11 xylanase BaxA | *Bacillus amyloliquefaciens* | AAZ17388.1 | – | S138T | 6 | 5 | 6.64 → 6.64 |
| 17 | Endo-1,4-$\beta$-xylanase | *Bacillus circulans* | P09850 | 1XNB | G34R | 5.5 | 6.5 | 6.27 → 6.68 |
| 17 | Endo-1,4-$\beta$-xylanase | *Bacillus circulans* | P09850 | 1XNB | Q175R | 5.5 | 7 | 6.27 → 6.8 |
| 17 | Endo-1,4-$\beta$-xylanase | *Bacillus circulans* | P09850 | 1XNB | Q7D | 5.5 | 5 | 6.27 → 6.12 |
| 17 | Endo-1,4-$\beta$-xylanase | *Bacillus circulans* | P09850 | 1XNB | V82D | 5.5 | 5 | 6.27 → 6.12 |
| 18 | Alkaline protease BgAP | *Bacillus gibsonii* | GN111900.1 | – | N253D | 11 | 10 | 9.28 → 9.24 |
| 18 | Alkaline protease BgAP | *Bacillus gibsonii* | GN111900.1 | – | Q256E | 11 | 10 | 9.28 → 9.24 |
| 18 | Alkaline protease BgAP | *Bacillus gibsonii* | GN111900.1 | – | N253D, Q256E | 11 | 10 | 9.28 → 9.23 |
| 19 | Endo-1,4-$\beta$-xylanase XYNII | *Trichoderma reesei* | P36217 | 1ENX | S186R, N67R, T26R, Q34R (ST4) | 5-5.5 | 6 | 7.16 → 7.2 |
| 19 | Endo-1,4-$\beta$-xylanase XYNII | *Trichoderma reesei* | P36217 | 1ENX | S186R, N67R, T26R, Q34R, S40R (ST5) | 5-5.5 | 6-6.5 | 7.16 → 7.23 |
| 19 | Endo-1,4-$\beta$-xylanase XYNII | *Trichoderma reesei* | P36217 | 1ENX | S186R, N67R, T26R, Q34R, N69R (ST6) | 5-5.5 | 6-6.5 | 7.16 → 7.23 |
| 20 | Acidic $\beta$-glucuronidase | *Aspergillus oryzae Li-3* | A7XS03 | 5C70 | E124R, E135R | 4.5 | 6.5 | 7.4 → 7.8 |
| 20 | Acidic $\beta$-glucuronidase | *Aspergillus oryzae Li-3* | A7XS03 | 5C70 | E150R, D200R | 4.5 | 6.5 | 7.4 → 7.8 |
| 20 | Acidic $\beta$-glucuronidase | *Aspergillus oryzae Li-3* | A7XS03 | 5C70 | E150R, D200R, E79R | 4.5 | 6.5 | 7.4 → 7.8 |
| 20 | Acidic $\beta$-glucuronidase | *Aspergillus oryzae Li-3* | A7XS03 | 5C70 | E150R, D200R, E124R | 4.5 | 6.5 | 7.4 → 7.8 |
| 20 | Acidic $\beta$-glucuronidase | *Aspergillus oryzae Li-3* | A7XS03 | 5C70 | E150R, D200R, E135R | 4.5 | 6.5 | 7.4 → 7.8 |
| 20 | Acidic $\beta$-glucuronidase | *Aspergillus oryzae Li-3* | A7XS03 | 5C70 | E150R, D200R, E124R, E79R | 4.5 | 6.5 | 7.4 → 7.8 |
| 20 | Acidic $\beta$-glucuronidase | *Aspergillus oryzae Li-3* | A7XS03 | 5C70 | E150R, D200R, E135R, E124R, E79R | 4.5 | 6.5 | 7.4 → 7.8 |

#### 3. Supplementary Figures

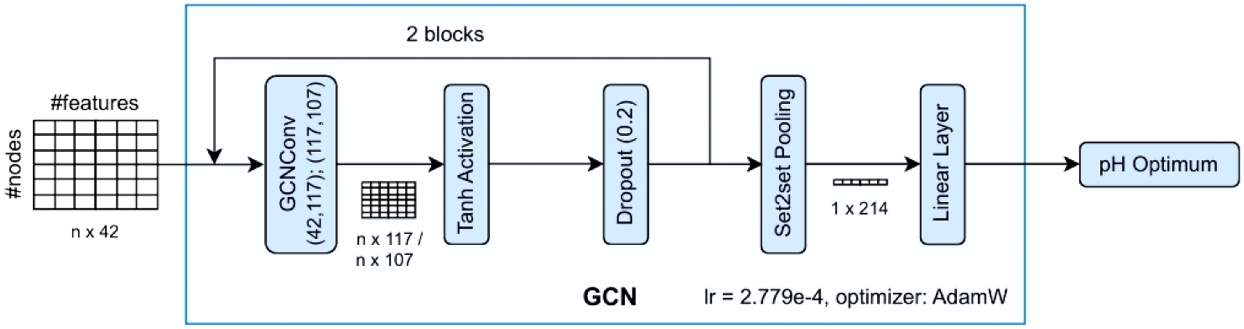


**Figure S1. Architecture of the optimal graph convolutional network (GCN).**

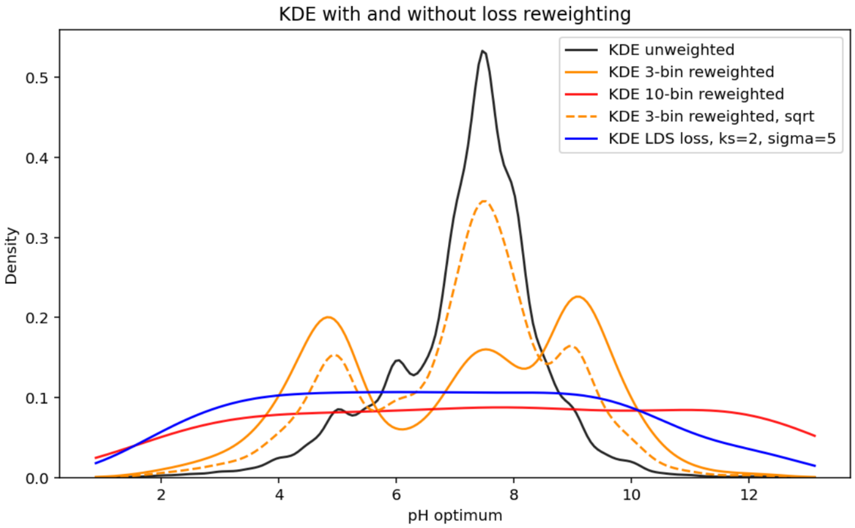


**Figure S2. Kernel Density Estimation (KDE) curves demonstrating the effect of various loss reweighting methods.**

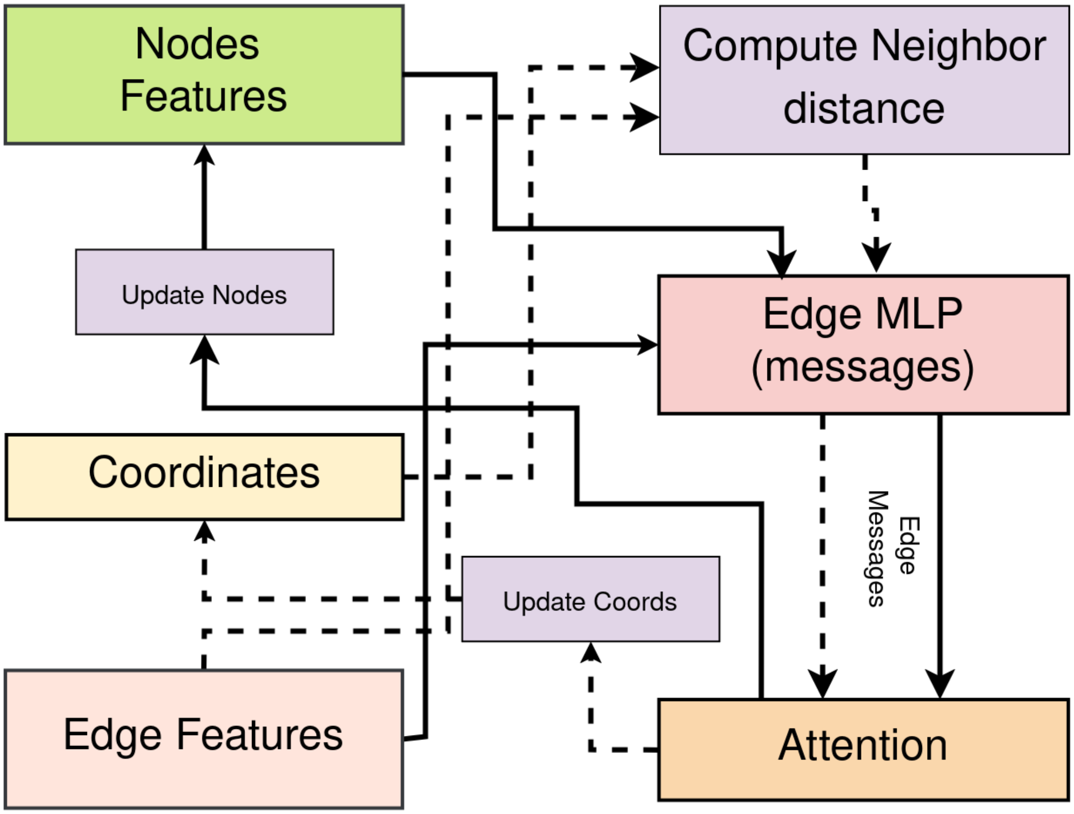


**Figure S3. Structure of a single EGCL layer with edge-level attention.** Node features and atomic coordinates are used to compute pairwise messages via an edge MLP. An attention module assigns a learnable weight to each directed interaction, modulating message strength before aggregation. The attention-weighted messages are then used to update both node features and atomic coordinates while preserving $E(n)$-equivariance.

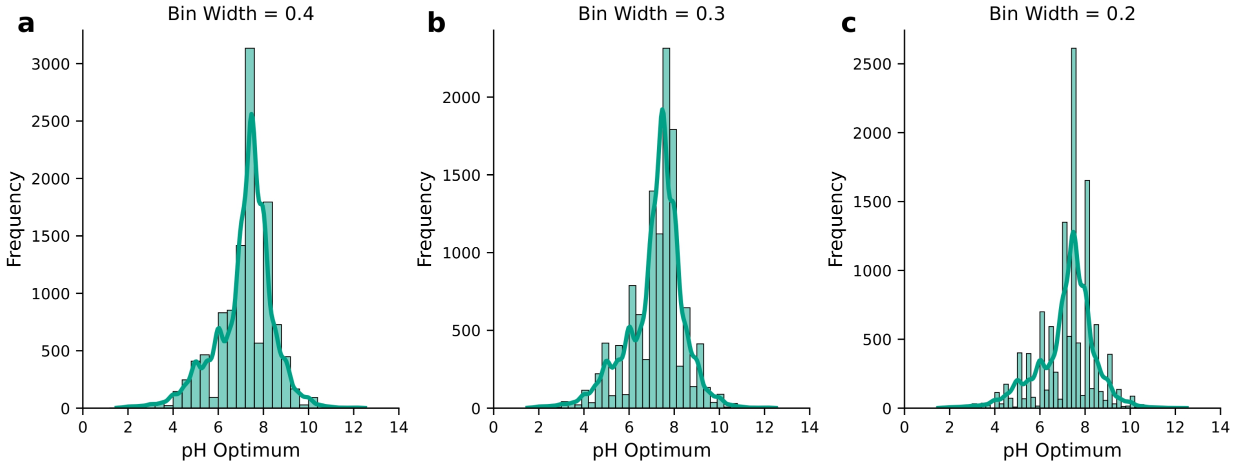


**Figure S4. Histograms of catalytic pH optima in EnzyBase12k across varying bin widths (0.4, 0.3, and 0.2 pH units).** Decreasing bin width highlights dataset imbalance, particularly the scarcity of data below pH 4 and above pH 10. Specific intervals (e.g., pH 6.1–6.5) are notably underrepresented compared to enriched regions (e.g., pH 6.9–7.1).

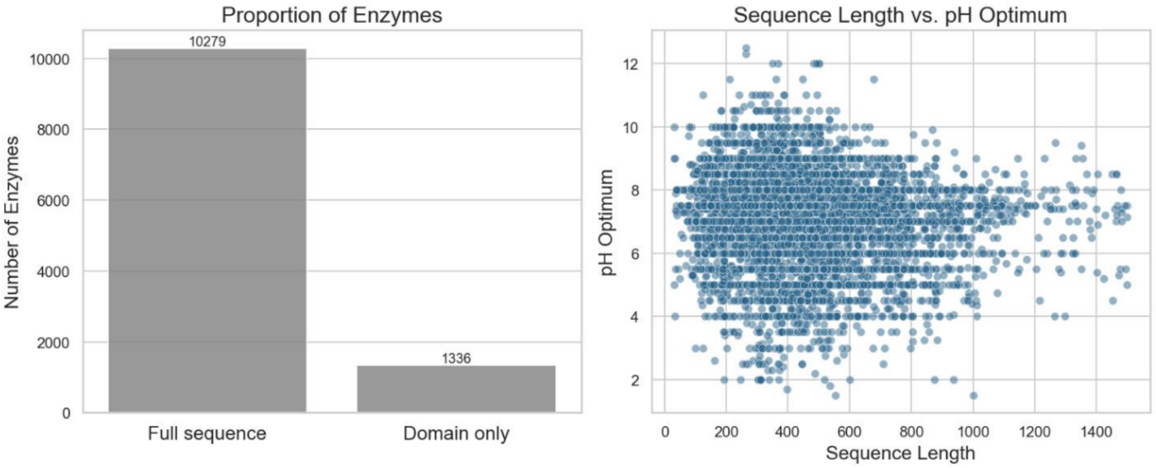


**Figure S5. Correlation analysis of domain-only proportions and sequence lengths with pH optima.** This figure illustrates the relationship between structural domain properties and the catalytic pH optima.

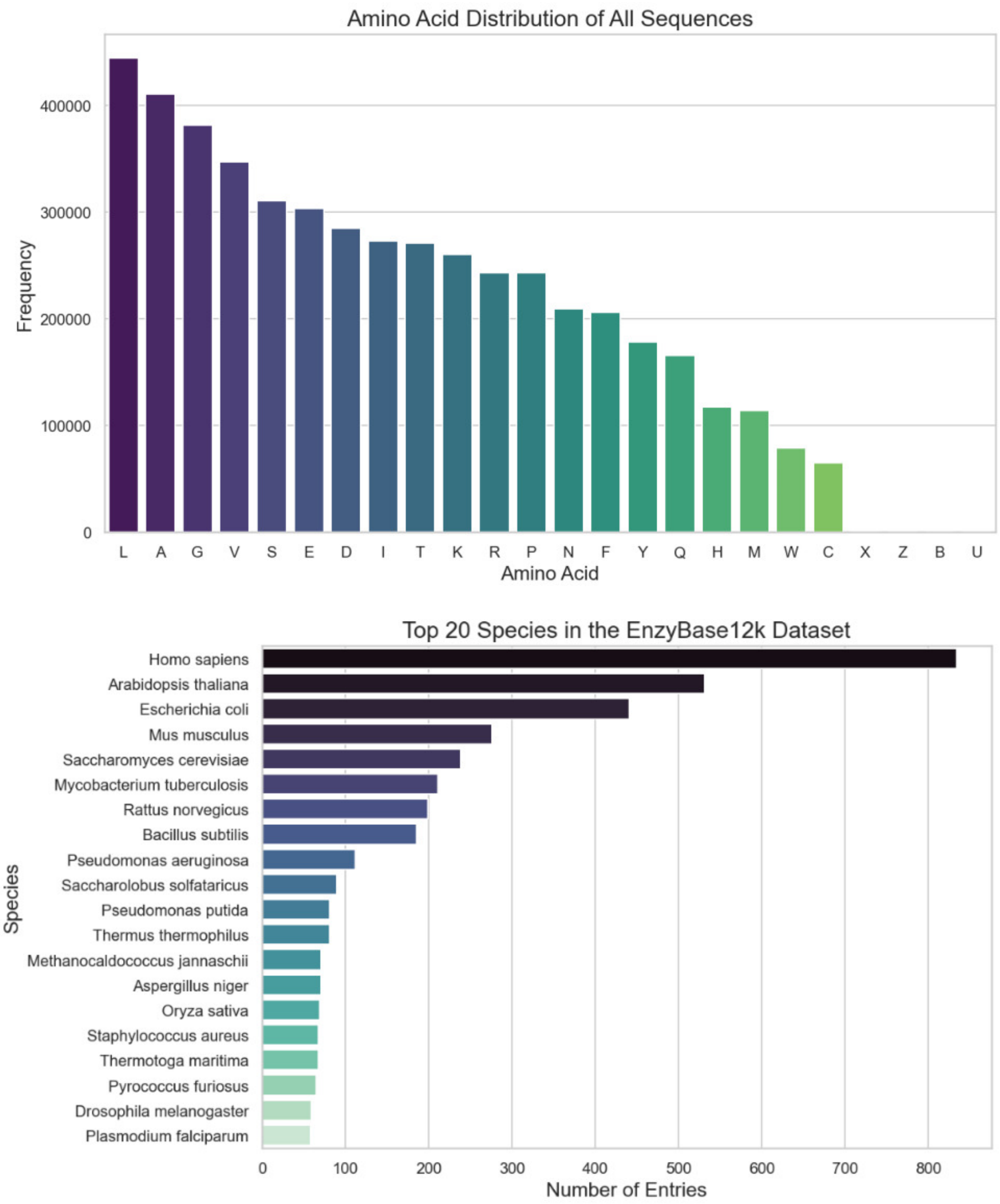


**Figure S6. Dataset composition statistics.** (Top) Amino acid distribution across all EnzyBase12k sequences. (Bottom) Frequency of the top 20 species represented in the EnzyBase12k dataset.

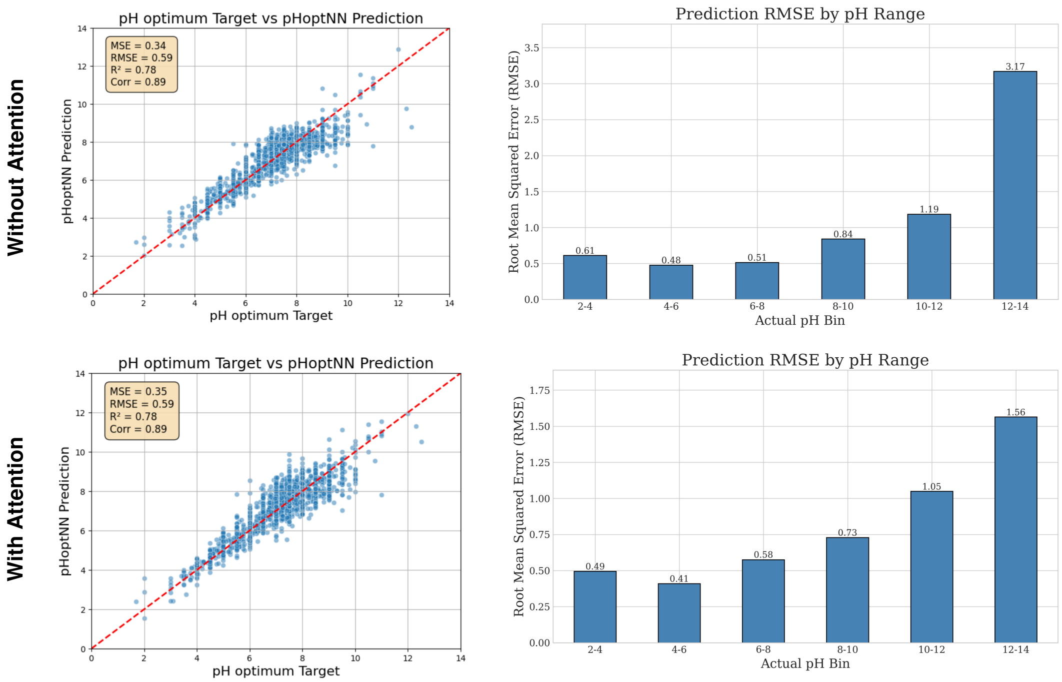


**Figure S7. Performance comparison of attention mechanisms.** (Top) Model predictions without attention. (Bottom) Model predictions with attention. While overall MSE and RMSE are comparable, the attention-based model demonstrates superior predictive performance in the alkaline pH range.

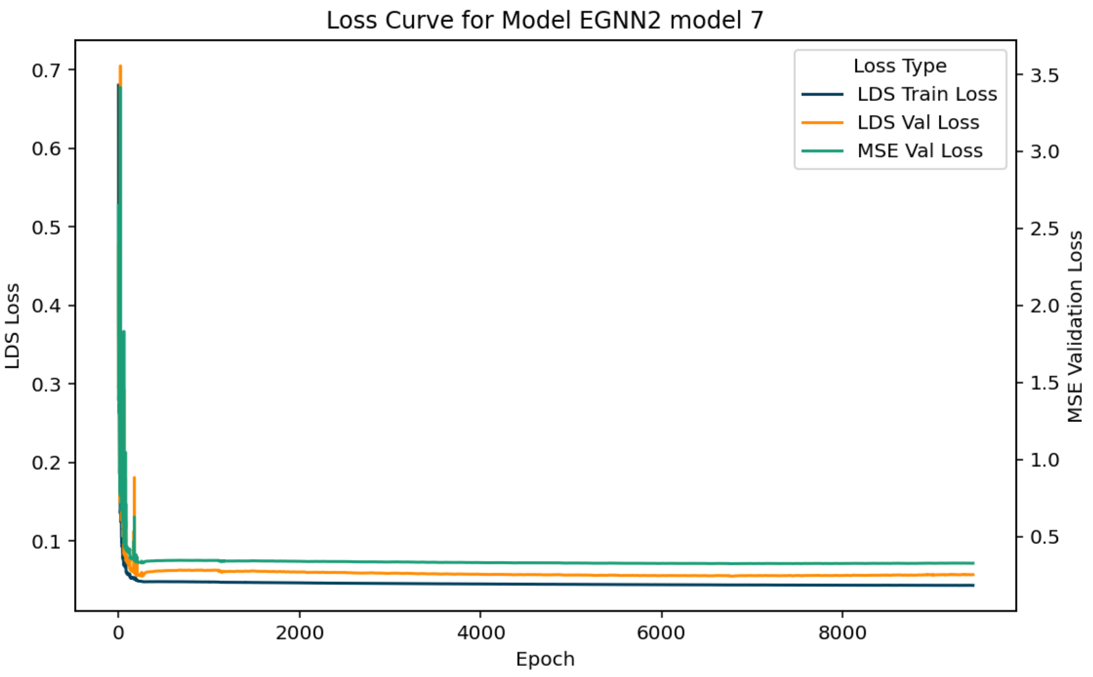


**(a)** Training and validation loss curves without early stopping.

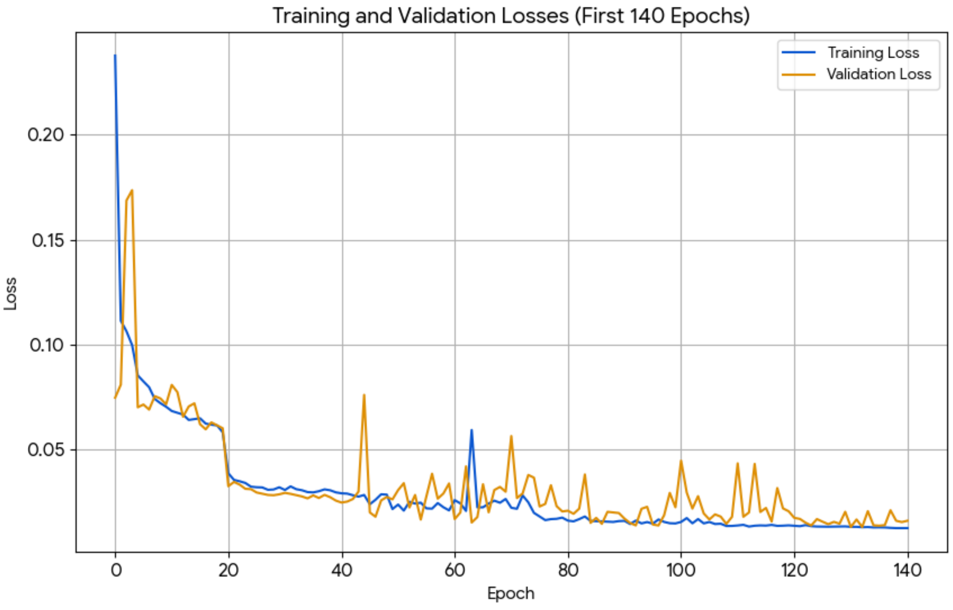


**(b)** Training and validation loss curves with early stopping (initial 140 epochs).

**Figure S8. Training dynamics of optimal models.** Comparison of loss trajectories under different training regimens.

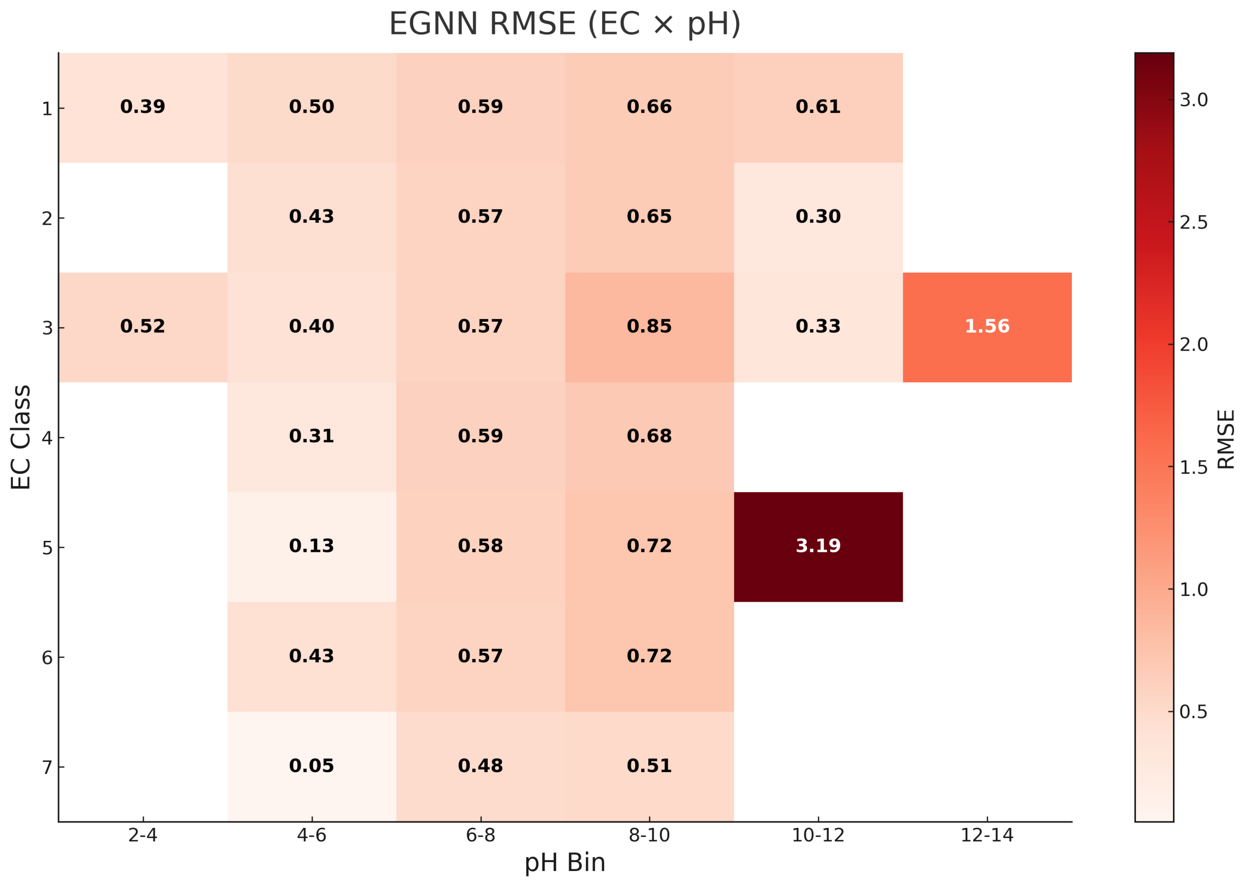


**Figure S9. RMSE heatmap by EC class and pH bin.** The model demonstrates strong performance across most categories. High RMSE values in the pH 12–14 range and pH 10–12 (EC class 5) are attributed to data sparsity in these regions.

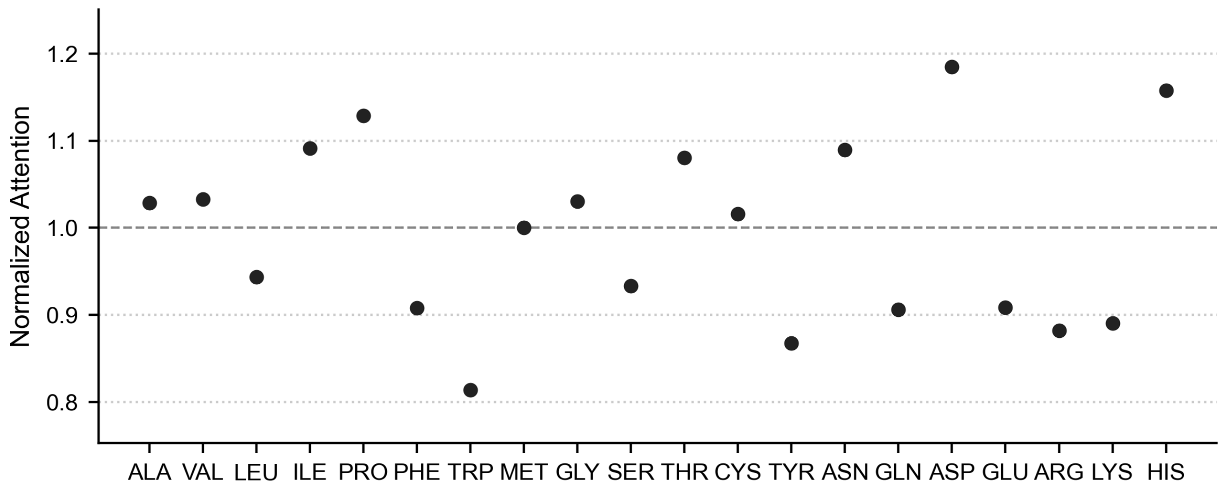


**Figure S10. Per-amino acid normalized attention weights.** Mean normalized attention weights for individual amino acid types are shown. Atom-level attention scores were aggregated to residues and normalized within each protein such that the mean attention across all residues equals 1.0 (dashed line), followed by averaging across proteins.

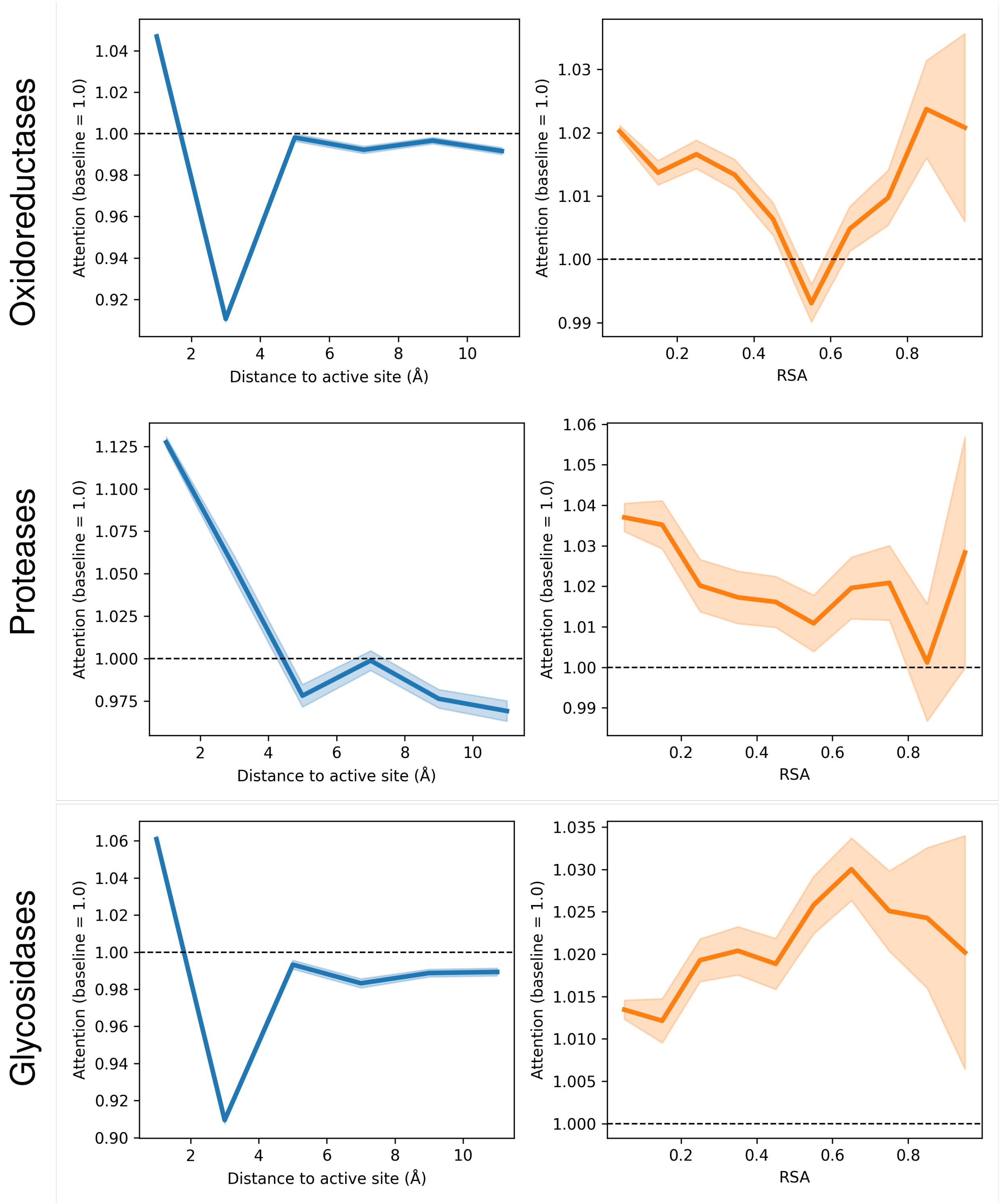


**Figure S11. Interpretability analysis of the pHoptNN attention mechanism across three distinct enzyme classes.** The learned attention weights were analyzed for Oxidoreductases, Proteases, and Glycosidases to assess feature importance. **Left Column:** Average attention scores plotted against the Euclidean distance from the catalytic active site (Å). The model demonstrates a strong localized focus, consistently assigning high significance (attention $> 1.0$) to residues within the immediate vicinity of the active site, with attention decaying below baseline beyond an 8 Å radius. **Right Column:** Attention scores relative to Relative Solvent Accessibility (RSA). The network generally places higher weight on highly solvent-exposed residues (RSA $\rightarrow 1.0$), reflecting the biophysical importance of surface electrostatics in determining pH optima. Shaded regions represent the standard deviation/confidence interval, and the horizontal dashed line indicates the baseline attention score of 1.0.

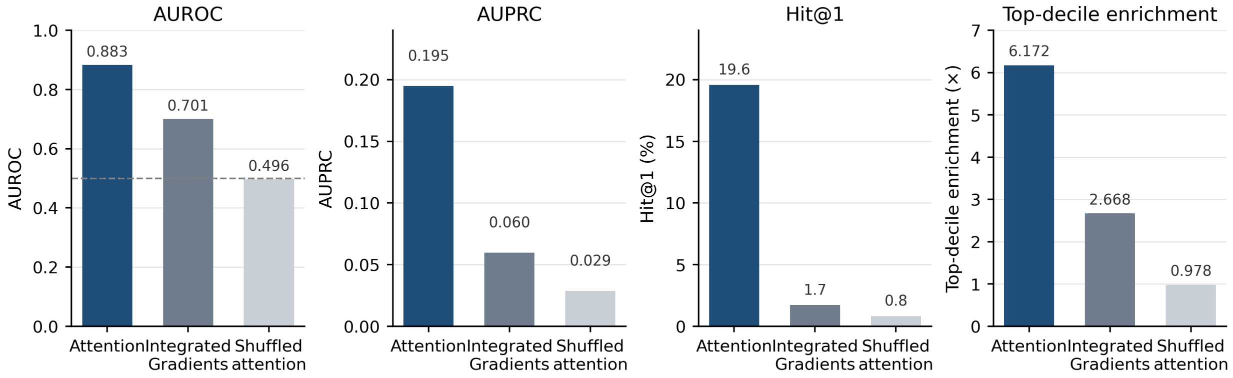


**Figure S12. Benchmark of residue-level attribution methods for catalytic site enrichment.** Comparison of the selected attention-based extraction strategy, Integrated Gradients, and shuffled attention for identifying annotated catalytic residues. The attention-based approach shows a consistent tendency toward higher enrichment across AUROC, AUPRC, Hit@1, and top-decile enrichment metrics relative to the baselines. These results suggest that the model-derived residue rankings are non-random and exhibit preferential association with functional residues, although they should not be interpreted as precise identification of catalytic sites.
